# Assembly of Long Error-Prone Reads Using de Bruijn Graphs

**DOI:** 10.1101/048413

**Authors:** Yu Lin, Jeffrey Yuan, Mikhail Kolmogorov, Max W. Shen, Pavel A. Pevzner

## Abstract

The recent breakthroughs in assembling long error-prone reads (such as reads generated by Single Molecule Real Time technology) were based on the overlap-layout-consensus approach and did not utilize the strengths of the alternative de Bruijn graph approach to genome assembly. Moreover, these studies often assume that applications of the de Bruijn graph approach are limited to short and accurate reads and that the overlap-layout-consensus approach is the only practical paradigm for assembling long error-prone reads. Below we show how to generalize de Bruijn graphs to assemble long error-prone reads and describe the ABruijn assembler, which results in more accurate genome reconstructions than the existing state-of-the-art algorithms.

## 1 Introduction

When the first reads generated using Single Molecule Real Time (SMRT) sequencing technology were made available [18], most researchers were skeptical about the ability of existing algorithms to generate high-quality assemblies from error-prone SMRT reads. Roberts et al., 2013 [51] even referred to this widespread skepticism as the “error myth” and argued that new assemblers for error-prone reads need to be developed to debunk this myth. Indeed, the key challenge for the success of SMRT and other recently emerged long reads technologies lies in the development of algorithms for assembling genomes from inaccurate reads.

The pioneer in long reads technologies, Pacific Biosciences, now produces accurate assemblies from error-prone SMRT reads [7,16]. Goodwin et al. [19] and Loman et al. [37] demonstrated that high-quality assemblies can be obtained from even less accurate Oxford Nanopore reads. Advances in assembly and mapping of long error-prone reads recently resulted in accurate assemblies of various genomes [28, 29, 31], reconstruction of complex regions of the human genome [15,22], and resolving complex tandem repeats [60]. However, as illustrated in Booher et al., 2015 [10], the problem of assembling long error-prone reads is far from being resolved even in the case of relatively short bacterial genomes.

All previous studies of SMRT assemblies were based on the *overlap-layout-consensus (OLC)* approach [26] or similar *string graph* approach [40], which require an all-against-all comparison of reads [39] and remain computationally challenging (see [23,33,44] for a discussion of *pros* and *cons* of this approach).

Moreover, there is an implicit assumption that the de Bruijn graph approach, which dominated genome assembly in the last decade, is inapplicable to assembling long reads. This is a misunderstanding since the de Bruijn graph approach, as well as its variation called the *A-Bruijn graph* approach, was developed to assemble rather long Sanger reads [45].

There is also a misunderstanding that the de Bruijn graph approach can only assemble highly accurate reads and fails while assembling error-prone SMRT reads, yet another “error myth” that we debunk in this paper. While this is true for the original de Bruijn graph approach to assembly [23, 44], the A-Bruijn graph approach was originally designed to assemble inaccurate reads as long as *any* similarities between reads can be reliably identified. Moreover, A-Bruijn graphs have proven to be useful even for assembling mass spectra, which represent highly inaccurate fingerprints of amino acid sequences of peptides [4, 5]. This A-Bruijn graph approach has turned the time-consuming sequencing of intact antibodies into a routine task [21, 59]. However, while A-Bruijn graphs have proven to be useful in assembling Sanger reads and mass spectra, the question of how to apply A-Bruijn graphs for assembling SMRT reads remains open.

De Bruijn graphs are a key algorithmic technique in genome assembly [23, 9, 12, 56, 61, 6]. In addition, de Bruijn graphs have been used for Sequencing by Hybridization [43], repeat classification [45], de novo protein sequencing [4,5,21], synteny block construction [38,46], multiple sequence alignment [48], genotyping [25], and immunoglobulin classification [13]. A-Bruijn graphs are even more general than de Bruijn graphs, e.g., they include *breakpoint graphs*, the workhorse of genome-rearrangement studies [42,35].

However, as discussed in [34], the original definition of a de Bruijn graph is far from being optimal for the challenges posed by the assembly problem. Below, we describe the concept of an A-Bruijn graph [45], introduce the ABruijn assembler for SMRT reads (including reads generated using Oxford Nanopore technology), and demonstrate that it generates accurate genome reconstructions.

## 2 Assembling long error-prone reads

Below we describe how ABruijn assembles long and error-prone reads into an error-prone *draft* genome. The challenge of assembling long error-prone reads. Booher et al., 2015 [10] recently sequenced important plant pathogens BLS256 and PXO99A representing *Xanthomonas* strains, and revealed the striking plasticity of *tal* genes which play key role in *Xanthomonas* infections. Each *tal* gene encodes a *TAL protein* that has a large domain formed by nearly identical *TAL repeats* (each TAL repeat has length ≈ 35 aa). Since variations in *tal* genes and TAL repeats are important for understanding the pathogenicity of various *Xanthomonas* strains, massive sequencing of these strains is an important task that may enable the development of novel plant disease control measures [55]. However, assembling *Xanthomonas* genomes using SMRT reads (let alone, short reads) remains challenging.

Depending on the strain, *Xanthomonas* genomes may harbor as many as 24 *tal* genes with each *tal* gene encoding 17 TAL repeats on average (some *tal* genes encode over 30 TAL repeats). These repeats render *tal* genes nearly impossible to assemble using short read technologies. Moreover, as Booher et al., 2015 [10] described, the existing SMRT assemblers also face challenges assembling *Xanthomonas* genomes, e.g., HGAP 2.0 failed to assemble BLS256. The assembly of BLS256 and PXO99A datasets is particularly challenging since these genomes have an unusually large numbers of *tal* genes (28 and 19, respectively).

### From de Bruijn graphs to A-Bruijn graphs

In the A-Bruijn graph framework, the classical de Bruijn graph *DB*(*String*, *k*) of a string *String* is defined as follows. Let *Path*(*String*, *k*) be a path consisting of |*String*| – *k* + 1 edges, where the *i*-th edge of this path is labeled by the *i*-th *k*-mer in *String* and the *i*-th vertex of the path is labeled by the *i*-th (*k*-1)-mer in *String*. The de Bruijn graph *DB* (*String*, *k*) is formed by gluing identically labeled vertices in *Path*(*String*, *k*) (Figure 1). Note that this somewhat unusual definition results in exactly the same de Bruijn graph as the standard definition (see [17] for details).

**Fig. 1.**
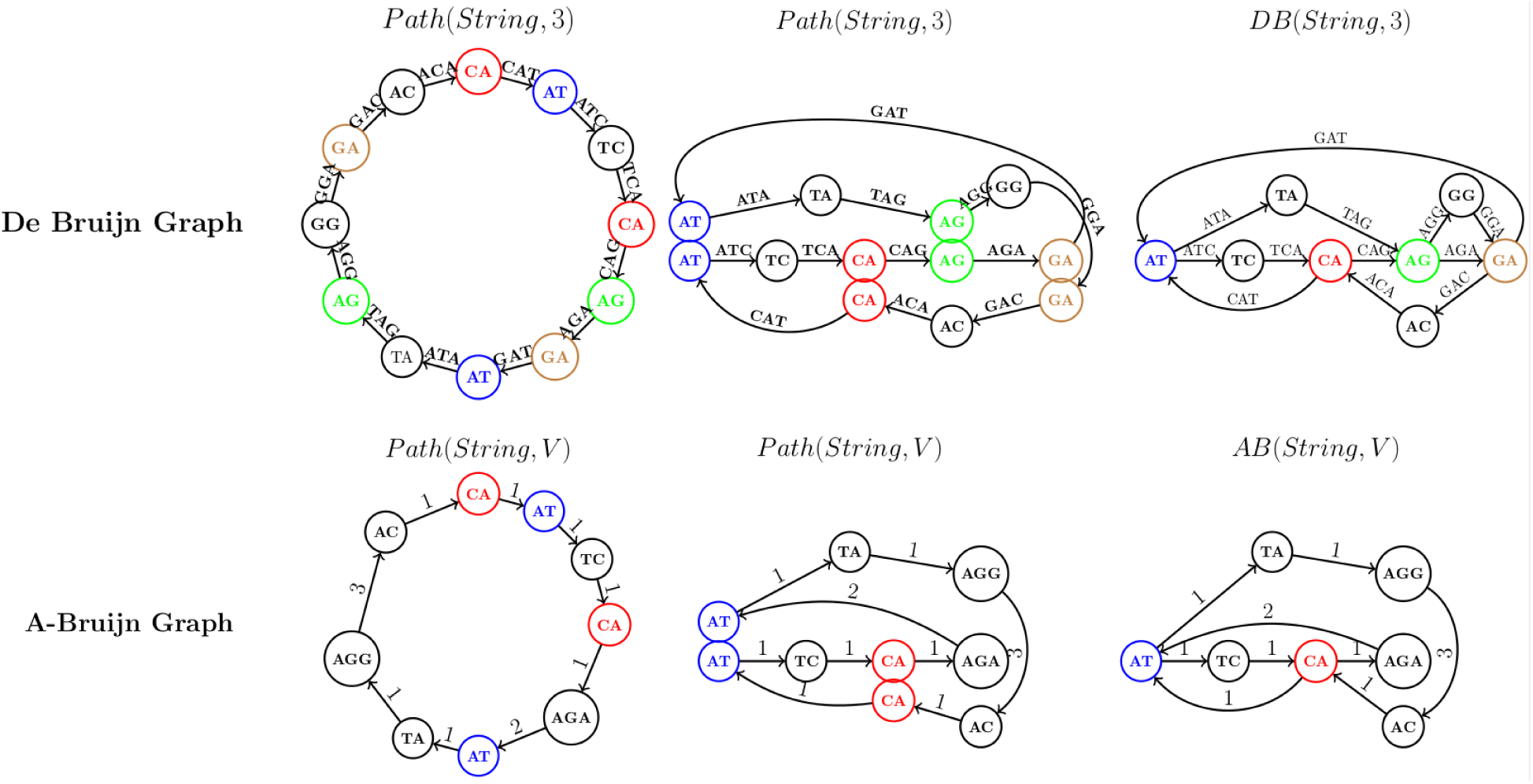
Constructing the de Bruijn graph (top) and the A-Bruijn graph (bottom) for a circular String=CATCAGATAGGA. (Top) From *Path*(*String*, 3) to *DB*(*String*, 3). (Bottom) From *Path*(*String*, *V*) to *AB*(*String*, *V*) for *V* = {CA, AT, TC, AGA, TA, AGG, AC}. The figure illustrates the process of bringing the vertices with the same label closer to each other to eventually glue them into a single vertex (middle column).

We now consider an *arbitrary* substring-free set of strings *V* (which we refer to as a set of *solid strings*), where no string in *V* is a substring of another one in *V*. The set *V* consists of words (of any length) and the new concept *Path*(*String*, *V*) is defined as a path through all words from *V* appearing in *String* (in order) as shown in Figure 1, bottom. We further assign integer *shift*(*v*, *w*) to the edge (*v*, *w*) in this path to denote the difference between the positions of *v* and *w* in *String* (i.e., the number of symbols between the start of *v* and the start of *w* in *String*). Afterwards, we glue identically labeled vertices as before to construct the A-Bruijn graph *AB* (*String*, *V*) as shown in Figure 1, bottom. Clearly, *DB* (*String*, *k*) is identical to *AB*(*String*, *Σ*^*k*–1^), where *Σ*^*k*–1^ stands for the set of all (*k*-1)-mers in alphabet *Σ*.

The definition of *AB* (*String*, *V*) naturally generalizes to *AB*(*Reads*, *V*) by constructing a path for each read and further gluing all identically labeled vertices in all paths. Since an Eulerian path in *AB* (*Reads*, *V*) spells out the genome [45], it appears that the only thing needed to apply the A-Bruijn graph concept to SMRT reads is to select an appropriate set of solid strings *V* and to construct the graph *AB*(*Reads*, *V*). Below we illustrate that this question is not as simple as it may appear and describe how it is addressed in the ABruijn assembler.

### Selecting solid strings for constructing A-Bruijn graphs

Different approaches to selecting solid strings affect the complexity of the resulting A-Bruijn graph and may either enable further assembly using the A-Bruijn graph or make it impractical. For example, when the set of solid strings *V* = *Σ*^*k*–1^ consists of all (*k*-1)-mers, *AB*(*Reads*, *Σ*^*k*–1^) may be either too tangled (if *k* is small) or too fragmented (if *k* is large).

While this is true for both short Illumina reads and long SMRT reads, there is a key difference between these two technologies with respect to their resulting A-Bruijn graphs. In the case of Illumina reads, there exists a range of values *k* so that one can apply various *graph simplification* procedures (e.g., *bubble* and *tip* removal [45,61]) to enable further analysis of the resulting graph. However, these graph simplification procedures were developed for the case when the error rate in the reads does not exceed 1% (like in the case of Illumina reads) and fail in the case of SMRT reads, with the error rate exceeding 10%. This complication led to the widespread opinion that the de Bruijn approach is not applicable to SMRT reads.

We argue that *k*-mers that frequently appear in reads (for sufficiently large *k*) are good candidates for the set of solid strings and define a (*k*,*t*)-*mer* as a *k*-mer that appears at least *t* times in the set *Reads*. We classify a *k*-mer as *genomic* if it appears in the genome and *non-genomic* otherwise. We further classify a *k*-mer as *unique (repeated)* if it appears once (multiple times) in the genome. Given a *k*-mer *a*, we define its *frequency* as the number of times this *k*-mer appears in reads. Figure 2 shows the histogram of the number of unique/repeated/non-genomic 15-mers with given frequencies for the *E. coli* SMRT dataset described in the “Datasets” section (referred to as ECOLI). As Figure 2 illustrates, the lion’s share of 15-mers with frequencies from 8 to 24 are unique 15-mers in the *E. coli* genome. Since non-genomic and repeated genomic *k*-mers complicate the analysis of the A-Bruijn graph [34], we remove all *k*-mers with frequencies exceeding *c* × *t* from the set of (*k*, *t*)-mers used as solid strings when constructing the A-Bruijn graph (the default value of the parameter *c* is 3).

**Fig. 2.**
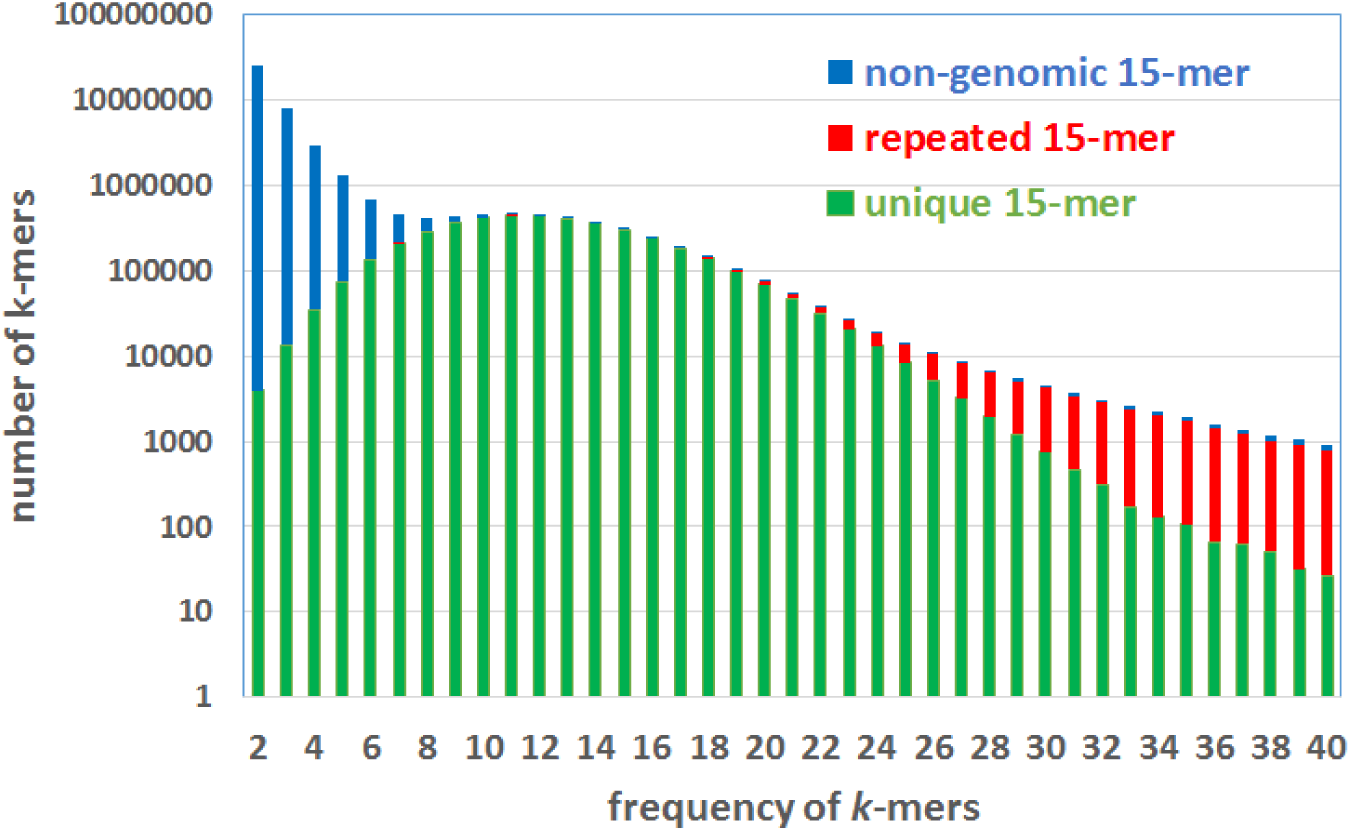
The histogram of the number of 15-mers with given frequencies for ECOLI dataset. The bars for unique/repeated/non-genomic 15-mers for the *E. coli* genome are stacked and shown in green/red/blue. ABruijn defines solid strings as all 15-mers with frequencies from 8 to 24.

For a typical bacterial SMRT assembly project with coverage 50X, ABruijn assembler uses *k* = 15 and *t* = 8 as the default choice. While larger values of *k* (typical for short read assemblies) also produce high-quality SMRT assemblies, we found that selecting smaller rather than larger *k* results in slightly better performance.

### Finding the genomic path in an A-Bruijn graph

After constructing an A-Bruijn graph, one faces the problem of finding a path in this graph that corresponds to traversing the genome (i.e., the *genomic path*) and correcting errors in the sequence spelled by this path. Since the SMRT reads are merely paths in the A-Bruijn graph, one can use the *path extension* paradigm [9,47, 58] to derive the genomic path from these (shorter) *read-paths*. exSPAnder [47] is a module of the SPAdes assembler [6] that finds a genomic path in the assembly graph constructed from short reads based either on read-pair paths or on SMRT read-paths like in hybridSPAdes [1]. Recent studies of bacterial plankton [30], antibiotics resistance [2], and genome rearangements [49] demonstrated that hybridSPades works well even for co-assembly with less accurate nanopore reads.

Below we sketch the hybridSPAdes algorithm for co-assembling short and long reads [1] and show how to modify the path extension paradigm to arrive at the ABruijn algorithm.

### hybridSPades

Given a set of paths *Paths* in a directed graph *Graph* and a parameter *minSupport*, we say that an edge in *Graph* is Paths-supported if it is traversed by at least *minSupport* paths in *Paths*. A set *Paths* is *consistent* (with respect to a parameter *minSupport*) if the set of all Paths-supported edges forms a single directed path in *Graph*. We further refer to this path as *ConsensusPath*(*Paths*, *minSupport*). The intuition for the notion of the consistent (inconsistent) set of paths is that they are sampled, with the exception of a relatively small number of chimeric paths, from a single segment (multiple segments) of the genomic path (see [1]).

Given two paths *P* and *P*′ in a weighted graph, we say that *P*′ *overlaps* with *P* if a sufficiently long suffix of *P* (of total weight at least *minOverlap*) coincides with a prefix of *P*′ and *P* does not contain the entire path *P*′ as a subpath. Given a path *P* and a set of paths *Paths*, we define *Paths*_*minOverlap*_(*P*) as the set of all paths in *Paths* that overlap with *P* (with respect to parameter *minOverlap*).

hybridSPAdes uses SPAdes to construct the de Bruijn graph from short reads and to further transform it into an *assembly graph* by removing bubbles and tips [6]. It further represents long reads as *read-paths* in the assembly graph and uses them for repeat resolution in the assembly graph. Our sketch of hybridSPAdes omits some details and deviates from the current implementation to make similarities with the A-Bruin graph approach more apparent, e.g., it only shows an algorithm for constructing a single contig.

**hybridSPAdes**(*ShortReads, LongReads, k, minSupport, minOverlap*)

construct the de Bruijn graph on k-mers from *ShortReads*

transform the de Bruin graph into the assembly graph

*ReadPaths* ← the set of paths in the assembly graph corresponding to

all reads from *LongReads*

*InitialPath* ← an arbitrary read-path from *ReadPaths*

*GrowingPath* ← *InitialPath*

**while** forever

*OverlapPaths ← ReadPaths*_*minOverlap*_ (*GrowingPath*)

**if** the set *OverlapPaths* is consistent (wrt parameter *minSupport*)

**if** *ConsensusPath*(*OverlapPaths*, *minSupport*) contains *InitialPath*

**return** the string spelled by *GrowingPath* (as the complete genome)

**if** *ConsensusPath*(*OverlapPaths*, *minSupport*) overlaps with

*GrowingPath*

extend *GrowingPath* by *ConsensusPath*(*OverlapPaths*, *minSupport*)

**else**

**return** the string spelled by *GrowingPath* (as one of the contigs)

### From hybridSPAdes to longSPAdes

Using the concept of the A-Bruijn graph, a similar approach can be applied to assembling long reads only. The pseudocode of longSPAdes differs from the pseudocode of hybridSPAdes by only the top three lines shown below:

**longSPAdes**(*LongReads, k, t, minSupport, minOverlap*)

construct the A-Bruijn graph on (*k*, *t*)-mers from *LongReads*

transform the A-Bruin graph into the assembly graph

We note that longSPAdes constructs a path spelling out an error-prone *draft genome* that requires further error-correction. However, error-correction of a *draft genome* is faster than the error correction of *individual reads before assembly* in the OLC approach [7, 16, 19, 37].

While hybridSPAdes and longSPAdes are similar, longSPAdes is more difficult to implement since bubbles in the A-Bruijn graph of error-prone long reads are more complex than bubbles in the de Bruijn graph of accurate short reads (see SI1: “Additional details on the ABruijn algorithm” for an example of a bubble in an A-Bruijn graph). As a result, the existing graph simplification algorithms fail in the case of the A-Bruin graph of long error-prone reads. While it is possible to modify the existing graph simplification procedure for SMRT reads (to be described elsewhere), this paper focuses on a different approach that does not require graph simplification.

### From longSPAdes to ABruijn

Instead of finding a genomic path in the simplified A-Bruijn graph, ABruijn attempts to find a genomic path in the original A-Bruijn graph. This approach leads to an algorithmic challenge: while it is easy to decide whether two reads overlap given an assembly graph, it is not clear how to answer the same question in the context of the A-Bruijn graph. Note that while the ABruijn pseudocode below uses the same terms *overlapping* and *consistent* as longSPAdes, these notions are defined differently in the context of the A-Bruijn graph. The new notions (as well as parameters *jump* and *maxOverhang*) are described later in this paper.

**ABruijn**(*LongReads, k, t, minSupport, minOverlap, jump, maxOverhang*)

construct the A-Bruijn graph on (*k*, *t*)-mers from *LongReads*

*ReadPaths* ← the set of paths in the assembly graph corresponding to

all reads from *LongReads*

*InitialPath* ← an arbitrary read-path in the A-Bruijn graph

*GrowingPath* ← *InitialPath*

*ReadPath* ← *InitialPath*

**while forever**

*OverlapPaths* ← all paths in *ReadPaths* overlapping *ReadPath*

(wrt *minOverlap, jump* and *maxOverhang*)

**if** the set *OverlapPaths* is consistent (wrt parameter *minSupport*)

**if** *InitialPath* is a consistent path in *OverlapPaths*

**return** the string spelled by *GrowingPath* (as the complete genome)

*ConsensusPath* ← a most-consistent path in *OverlapPaths*

(wrt parameter *minSupport*)

extend *GrowingPath* by *ConsensusPath*

*ReadPath ← ConsensusPath*

**else**

**return** the string spelled by *GrowingPath* (as one of the contigs)

The constructed path in the A-Bruijn graph spells out an error-prone draft genome (or one of the draft contigs). ABruijn uses a new approach to error-correction that first builds yet another A-Bruijn graph of reads aligned to the draft genome (for simplicity, the pseudocode above describes construction of a single contig and does not cover the error-correction step).

We note that while the A-Bruijn graph constructed *from reads* is very complex, the A-Bruijn graph constructed *from reads aligned to the draft genome* is rather simple. While there are hundreds of thousands of bubbles in this graph, each bubble is very simple, making the error correction step fast and accurate.

### Common *jump*-subpaths

Given a path *P* in a weighted directed graph (weights correspond to shifts in the A-Bruijn graph), we refer to the distance *d*_*P*_(*v*, *w*) along path *P* between vertices *v* and *w* in this path (i.e., the sum of the weights of all edges in the path) as the *P-distance*. The *span* of a subpath of a path *P* is defined as the *P*-distance from the first to the last vertex of this subpath.

Given a parameter *jump*, a *jump-subpath* of *P* is a subsequence of vertices *v*_1_ … *v*_*t*_ in *P* such that *d*_*P*_(*v*_*i*_, *v*_*i*+1_) ≤ *jump* for all *i* from 1 to *t*-1. We define *Path*_*jump*_(*P*) as a *jump*-subpath with the maximum span out of all *jump*-subpaths of a path *P*. We further denote the *span* of this *jump*-subpath as |*Path*_*jump*_(*P*)|.

A sequence of vertices in a weighted directed graph is called a *common jump-subpath* of paths *P*_1_ and *P*_2_ if it is a *jump*-subpath of both *P*_1_ and *P*_2_. The *span* of a common *jump*-subpath of *P*_1_ and *P*_2_ is defined as its span with respect to path *P*_1_ (note that this definition is non-symmetric with respect to *P*_1_ and *P*_2_). We refer to a common *jump*-subpath of paths *P*_1_ and *P*_2_ with the maximum span as *Path*_*jump*_(*P*_1_, *P*_2_) (with ties broken arbitrarily).

Below we describe how the ABruijn assembler uses the notion of common *jump*-subpaths with maximum span to detect overlapping reads.

### Finding a common *jump*-subpath with maximum span

For the sake of simplicity, below we limit attention to the case when paths *P*_1_ and *P*_2_ traverse each of their shared vertices exactly once.

A vertex *w* is a *jump*-predecessor of a vertex *v* in a path *P* if *P* traverses *w* before traversing *v* and *d*_*P*_(*w*, *v*) ≤ *jump*. We define *P*(*v*) as the subpath of *P* from its first vertex to *v*. Given a vertex *v* shared between paths *P*_1_ and *P*_2_, we define *span*_*jump*_(*v*) as the largest span among all common jump-subpaths of paths *P*_1_(*v*) and *P*_2_(*v*) ending in *v*. The dynamic programming algorithm for finding a common *jump*-subpath with the maximum span is based on the following recurrence:

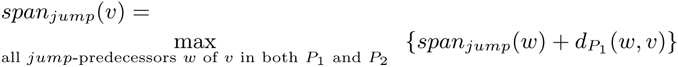

Note that finding a common jump-subpath with the maximum span becomes trivial when paths *P*_1_ and *P*_2_ traverse their shared vertices in the same order (more than half of all overlapping read-paths satisfy this condition). Note that, given a set of all paths sharing vertices with a path *P*, computing common jump-subpaths with maximum span with *P* for *all* of them can be done using a single scan of *P*. See SI1: “Additional details on the ABruijn algorithm” for a fast heuristic for finding a common jump-subpath with maximum span.

### Overlapping paths in A-Bruin graphs

We define the *right overhang* between paths *P*_1_ and *P*_2_ as the minimum of the distances from the last vertex in *Path*_*jump*_(*P*_1_, *P*_2_) to the ends of *P*_1_ and *P*_2_. Similarly, the *left overhang* between paths *P*_1_ and *P*_2_ is the minimum of the distances from the starts of *P*_1_ and *P*_2_ to the first vertex in *Path*_*jump*_(*P*_1_, *P*_2_).

Given parameters *jump, minOverlap* and *maxOverhang*, we say that paths *P*_1_ and *P*_2_ *overlap* if they share a common *jump*-subpath of span at least *minOverlap* and their right and left overhangs do not exceed *maxOverhang*. To decide whether two reads have arisen from two overlapping regions in the genome, ABruijn checks whether their corresponding read-paths *P*_1_ and *P*_2_ overlap (with respect to parameters *jump, minOverlap*, and *maxOverhang*). SI1: “Additional details on the ABruijn algorithm” describes how ABruijn detects chimeric reads. SI2: “Choice of parameters in the ABruijn algorithm” describes the range of parameters that work well for bacterial genome assembly.

### Consistent paths

Although it appears that the notion of overlapping paths allows us to implement the path extension paradigm for A-Bruijn graphs, there are two complications. First, while the algorithm for analyzing chimeric reads removes the lion’s share of chimeric reads, 0.5% of chimeric reads evade this algorithm and may end up in the set of overlapping read-paths extending *GrowingPath* in the ABruijn algorithm. Second, since some of the paths (i.e., reads) overlapping with *GrowingPath* may have large insertions and deletions, we want to exclude them when ABruijn selects a read-path during the path extension. SI1: “Additional details on the ABruijn algorithm” describes the concept of a *most-consistent* path which addresses these complications. Given a set of paths *Paths* overlapping with *ReadPath*, ABruijn selects a most-consistent path for extending *ReadPath*. Also, the simplified ABruijn pseudocode is limited to generating a single contig. In reality, after a contig is constructed, ABruijn maps all reads to this contig and uses the remaining reads to iteratively costruct other contigs.

## 3 Correcting errors in the draft genome

Below we describe how ABruijn corrects errors in the draft genome.

### Matching reads against the draft genome

ABruijn uses BLASR [14] to align all reads in the ECOLI dataset against the draft genome (92% of all reads align to the draft genome over at least a 5 kb segment). It further combines pairwise alignments of all reads to the draft genome into a multiple alignment *Alignment*. Since this alignment against the error-prone draft genome is rather inaccurate, we need to modify it into a different alignment that we will use for error correction.

Our goal now is to partition the multiple alignment of reads to the entire draft genome into hundreds of thousands of short segments (*mini-alignments*) and to error-correct each segment into the consensus string of the mini-alignment. The motivation for constructing mini-alignments is to enable accurate error-correction methods (such as a partial order alignment [32]) that are fast when applied to short segments of reads but become too slow in the case of long segments.

The task of constructing mini-alignments is not as simple as it may appear. For example, breaking this alignment into segments of fixed size will result in inaccurate consensus sequences since a region in a read aligned to a particular segment of the draft genome has not necesarily arisen from this segment, e.g., it may have arisen from a neighboring segment or from a different instance of a repeat. We thus search for a *good partition* of the draft genome that satisfies the following criteria: (i) most segments in the partition are short, so the algorithm for constructing the partial order alignment is fast, and (ii) with high probability, the region of each read aligned to a given segment in the partition represents a version (possibly error-prone) of *this* segment. Below we show how to address the challenge of constructing a good partition by building an A-Bruijn graph 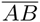 (*Alignment*) (Figure 3).

**Fig. 3.**
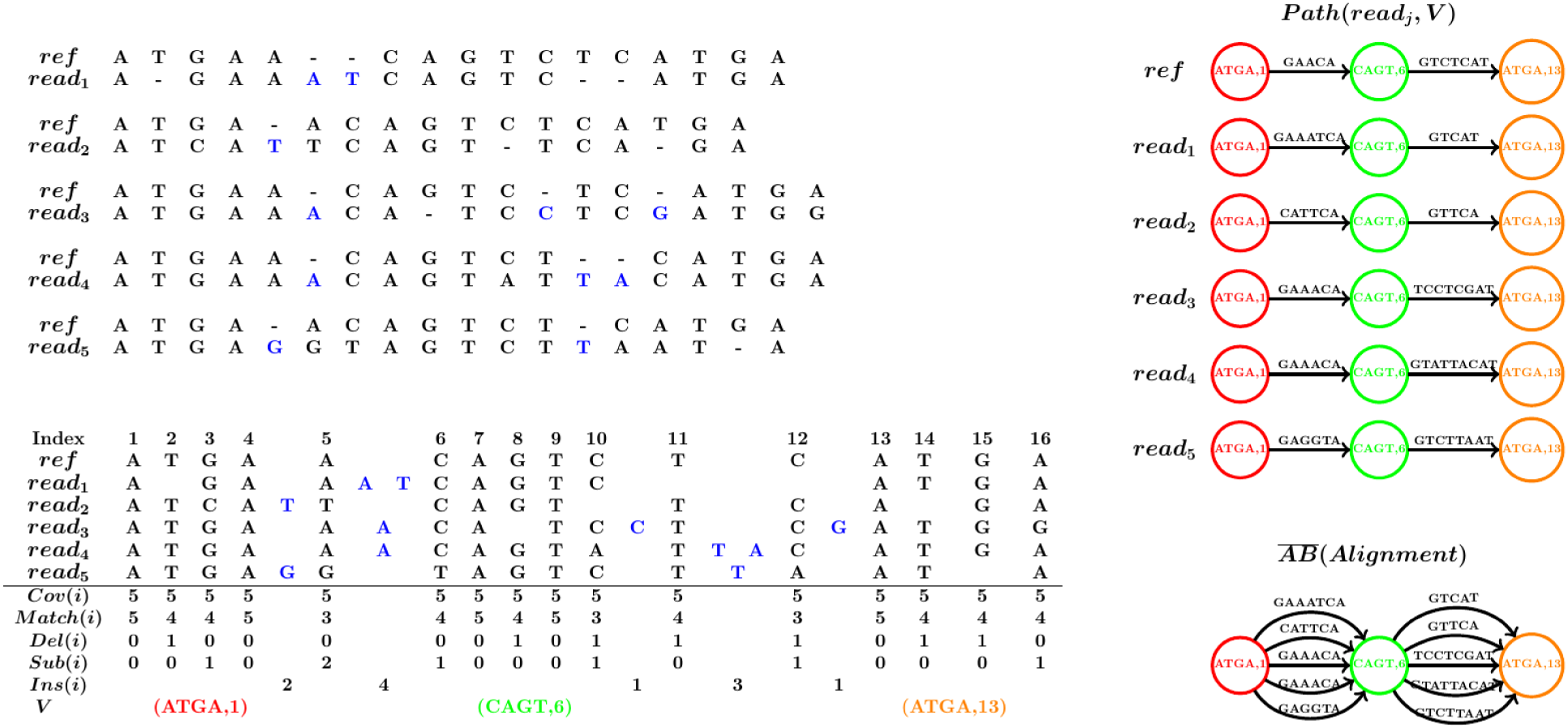
(Top Left) The pairwise alignments between a reference region *ref* in the draft genome and 5 reads *Reads* = {*read*_1_, *read*_2_, *read*_3_, *read*_4_, *read*_5_}. All inserted symbols in these reads with respect to the region *ref* are colored in blue. (Bottom Left) The multiple alignment *Alignment* constructed from the above pairwise alignments along with the values of *Cov*(*i*), *Match*(*i*), *Del*(*i*), *Sub*(*i*) and *Ins*(*i*). The last row shows the set *V* of (0.8, 0.2)-solid 4-mers. (Right) Constructing 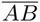 (*Alignment*), i.e., combining all paths *Path*(*read*_*j*_, *V*) into 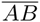(*Alignment*). Note that the 4-mer ATGA correspond to two different nodes with labels 1 and 13. The three bubble boundaries for this example are between positions 2 and 3, 7 and 8, and 14 and 15.

### Defining solid regions in the draft genome

We refer to a position (column) of the alignment with the space symbol “-” in the reference sequence as a *non-reference position* (*column*) and to all other positions as a *reference position* (*column*). We refer to the column in the multiple alignment containing the *i*-th position in a given region of the reference genome as the *i*-th column. As illustrated in Figure 3, the non-reference columns in the alignment are not numbered. The total number of reads covering a position *i* in the alignment is referred to as *Cov*(*i*).

A non-space symbol in a reference column of the alignment is classified as a *match (resp., substitution)* if it matches (resp., does not match) the reference symbol in this column. A space symbol in a reference column of the alignment is classified as a *deletion*. We refer to the number of matches, substitutions, and deletions in the *i*-th column of the alignment as *Match*(*i*), *Sub*(*i*), and *Del*(*i*), respectively. We refer to a non-space symbol in a non-reference column as an *insertion* and denote *Ins*(*i*) as the number of nucleotides in the non-reference columns flanked between the reference columns *i* and *i* + 1 (Figure 3).

For each reference position *i*, *Cov*(*i*) = *Match*(*i*) + *Sub*(*i*) + *Del*(*i*). We define the *match* rate, the *substitution* rate, the *deletion* rate, and the *insertion* rate at position *i* as *Match*(*i*)/*Cov*(*i*), *Sub*(*i*)/*Cov*(*i*), *Del*(*i*)/*Cov*(*i*), and *Ins*(*i*)/*Cov*(*i*), respectively.

Given an *l*-mer in a draft genome, we define its *local match rate* as the *minimum* match rate among the positions within this *l*-mer. We further define its *local insertion rate* as the *maximum* insertion rate among the positions within this *l*-mer.

An *l*-mer in the draft genome is called (*α*, *β*)-*solid* if its local match rate exceeds *α* and its local insertion rate does not exceed *β*. When *α* is large and *β* is small, (*α*, *β*)-solid *l*-mers typically represent the correct *l*-mers from the genome. SI3: “Additional details on constructing necklaces” describes how to use the draft genome to construct mini-alignments, demonstrates that (0.8,0.2)-solid *l*-mers in the draft genome are extremely accurate, and describes the choice of parameters for specifying (*α*, *β*)-solid *l*-mers that work well for assembly. The last row in Figure 3 (bottom left) shows all of the (0.8,0.2)-solid 4-mers (ATGA, CAGT, and ATGA) in the draft genome.

The contiguous sequence of (*α*, *β*)-solid l-mers forms a *solid region*. There are 1,146,866 (0.8,0.2)-solid 10-mers in the draft genome for the ECOLI dataset, which form 141,658 solid regions. Our goal now is to select a position (*landmark*) within each solid region and to form mini-alignments from the segments of reads spanning the intervals between two consecutive landmarks.

### Breaking the multiple alignment into mini-alignments

Since (*α*, *β*)-*solid* l-mers are very accurate (for appropriate choices of *α*, *β* and *l*), we use them to construct yet another A-Bruijn graph with much simpler bubbles. Since analyzing errors in homonucleotide runs is a difficult problem [16], we select landmarks outside homonucleotide runs (SI3: “Additional details on constructing necklaces” describes how ABruijn selects landmarks). ABruijn analyzes each mini-alignment and error-corrects each segment between consecutive landmarks (the average length of these segments is only ≈35 nucleotides). This procedure results in 135,417 mini-alignments.

### Constructing the A-Bruijn graph on solid regions in the draft genome

We label each solid region containing a landmark by its landmark position in *Alignment* and break each read into a sequence of segments aligned between consecutive landmarks. We further represent each read as a directed path through the vertices corresponding to the landmarks that it spans over. To construct the A-Bruijn graph 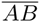 (*Alignment*), we glue all identically labeled vertices in the set of paths resulting from the reads (Figure 3 (right)).

Labeling vertices by their positions in the draft genome (rather than the sequences of solid regions) distinguishes identical solid regions from different regions of the genome and prevents excessive gluing of vertices in the A-Bruijn graph 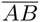(*Alignment*).

The edges between two consecutive landmarks (two vertices in the A-Bruijn graph) form a *necklace* consisting of segments from different reads that align to the region flanked by these landmarks. The SI3: “Additional details on constructing necklaces” describes how ABruijn constructs a consensus for each necklace (*necklace consensus*) and transforms the inaccurate draft genome for the ECOLI dataset into a rather accurate *pre-polished* genome with an error rate of only 0.1%.

We define the *length* of a necklace as the median length of its segments and classify a necklace as *long* if its length exceeds 100 bp. Although only 2,914 out of 135,417 necklaces constructed for the ECOLI dataset are long, their analysis takes the lion’s share of the running time for the error correction step in the ABruijn assembler. The SI3: “Additional details on constructing necklaces” describes how ABruijn reduces the number of long necklaces (from 2,914 to 135), at the expense of increasing the number of necklaces (from 135,417 to 283,909); this reduces the overall running time. Below we describe the algorithm to error-correct the pre-polished genome into a polished genome and to reduce the error rate from 0.1% to 0.0005% for ECOLI dataset (only 25 putative errors for the entire genome).

### A probabilistic model for necklace polishing

Each necklace contains read-segments *Segments* = {*seg*_1_, *seg*_2_, …, *seg*_*m*_}, and our goal is to find a consensus sequence *Consensus* maximizing *Pr*(*Segments*|*Consensus*) 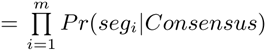, where *Pr*(*segi*|*Consensus*) is the probability of generating a segment *segi* from a consensus sequence *Consensus*. Given an alignment between a segment *seg*_*i*_ and a consensus *Consensus*, we define *Pr*(*seg*_*i*_|*Consensus*) as the product of all match, mismatch, insertion, and deletion rates for all positions in this alignment.

The match, mismatch, insertion, and deletion rates should be trained using an alignment of any set of reads (generated with the same technology) to any reference genome. Interestingly, the statistical parameters of the P6-C4 ECOLI dataset are nearly identical to the parameters of P5-C3 and even those of the older P4-C2 protocol for generating SMRT reads (SI5: “ Statistical analysis of errors in reads” describes statistical parameters for the P6-C4, P5-C3 and P4-C2 datasets).

Starting from the initial necklace consensus (see SI3: “Additional details on aconstructing necklaces”), ABruijn iteratively checks whether the consensus sequence for each necklace can be improved by introducing a single insertion, deletion or substitution. If there exists a mutation that increases *Pr(Segments\Consensus)*, we select the mutation that results in the maximum increase and iterate until convergence. We further output the final sequence as the error-corrected sequence of the necklace. As described in [16], this greedy strategy can be implemented efficiently since a mutation maximizing *Pr*(*Segments*|*Consensus*) among all possible mutated sequences can be found in a single run of the forward-backward dynamic programming algorithm for each sequence in *Segments*. The error rate after this step drops from 0.1% to 0.003%.

### Error-correcting homonucleotide runs

The probabilistic approach from [16] described above works well for most necklaces but its performance deteriorates when it faces the difficult problem of estimating the lengths of homonucleotide runs, which account for 46% of the *E. coli* genome (see discussion on *pulse merging* in [16]). We thus complement this approach with a *homonucleotide likelihood function* based on the statistics of homonucleotide runs. In contrast to previous approaches to error-correction of SMRT reads, this new likelihood function incorporates all corrupted versions of all homonucleotide runs across the training set of reads and reduces the error rate six-fold (from 0.003% to 0.0005%) compared to the standard likelihood approach.

To generate the statistics of homonucleotide runs for a given experimental protocol, we need an arbitrary set of reads (generated using this protocol) aligned against a training reference genome. For each homonucleotide run in the genome and each read spanning this run, we represent the aligned segment of this read simply as the set of its nucleotide counts. For example, if a run AAAAAAA in the genome is aligned against AATTACA in a read, we represent this read-segment as 4A1C2T. After collecting this information for all runs of AAAAAAA in the reference genome, we obtain the statistics for all read segments covering all instances of the homonucleotide run AAAAAAA (see the table in SI5: “ Statistical analysis of errors in reads”). We further use the frequencies in this table for computing the likelihood function as the product of these frequencies for all reads in each necklace (frequencies below a threshold 0.001 are ignored). Similar to the results in SI5: “ Statistical analysis of errors in reads”, the frequencies in the resulting table hardly change when one changes the dataset of reads, the reference genome, or even the SMRT protocol from P6-C4 to the older P5-C3. To decide on the length of a homonucleotide run, we simply select the length of the run that maximizes the likelihood function. For example, if *Segments*={5A, 6A, 6A, 7A, 6A1C}, *Pr*(*Segments*|6*A*)=0.155 × 0.473^2^ × 0.1 × 0.02 > *Pr*(*Segments*|7*A*)=0.049 × 0.154^2^ × 0.418 × 0.022 and we select AAAAAA over AAAAAAA as the necklace consensus.

While the described error-correcting approach results in a very low error rate even after a single iteration, ABruijn re-aligns all reads and error-corrects the pre-polished genome in an iterative fashion (three iterations by default).

## 4 Results

### Datasets

The *E. coli K12* SMRT dataset [27] (refered to as *ECOLI*) contains 10,277 reads with ≈55X coverage generated using the latest P6-C4 Pacific Biosciences technology (all reads are at least 20 kb long).

The *E. coli K12* Oxford Nanopore dataset [37] (refered to as ECOLI_*nano*_) contains 22,270 reads with ≈29X coverage (5,997 out of these 22,270 reads are at least 9 kb long). 99.6% of these 5,997 reads align to the *E. coli K12* reference genome over at least a 5 kb segment.

The BLS256 and PXO99A datasets were derived from two plant parasites *Xanthomonas oryzae* strains BLS256 (4,831,739 nucleotides) and PXO99A (5,240,075 nucleotides) previously assembled using Sanger reads [8, 53] and re-assembled using Pacific Biosciences P6-C4 reads in Booher et al., 2015 [10] (SRA accession numbers SRX502906 and SRX502899, respectively). The BLS256 and PXO99A datasets contains 21,996 and 17,577 long reads (longer than 14 kb), respectively.

### Benchmarking

ABruijn assembles the ECOLI, ECOLI_*nano*_, and BLS256 datasets into a single circular contig structurally concordant with the *E. coli* genome (see SI4: “Draft ECOLI assembly”). It also assembled the PXO99A dataset into a single circular contig structurally concordant with the PXO99A reference genome but, similarly to the initial assembly in Booher et al., 2015, it collapsed a 212 kb tandem repeat. Below we focus on assembling the ECOLI dataset and describe other assemblies in SI7: “Additional details on assembling Oxford Nanopore reads.” and SI8 “Assembling *Xanthomonas* genomes.”

Evaluating the accuracy of SMRT assemblies should be done with caution. For example, high-quality short-read assemblies often have error-rates on the order of 10^‒5^, which typically result in 50–100 errors per assembled genome [52]. Since assemblies of high-coverage SMRT datasets are often even more accurate than assemblies of short reads, short-read assemblies do not represent a gold standard for estimating the accuracy of SMRT assemblies. Moreover, as the ECOLI dataset reveals, the *E. coli K12* strain used for SMRT sequencing differs from the reference genome (see SI6: “Differences between the genome that gave rise to ECOLI dataset and the reference *E. coli K12* genome”). Thus, the standard benchmarking approach based on comparison with the reference genome [20] is not applicable to these assemblies.

We thus used the following approach to benchmark ABruijn and PBcR against the reference *E. coli K12* genome. There are 2906 and 2925 positions in *E. coli K12* genome where the reference sequence differs from ABruijn and PBcR, respectively. However, ABruijn or PBcR agree on 2871 of them, suggesting that these positions represent errors in the reference genome (or, more likely, mutations in *E. coli K12* as compared to the reference genome). We thus focused on the remaining positions where ABruijn and PBcR disagree with the reference strain and with each other. We further classify a position as an *ABruijn error* if the PBcR sequence at this position agrees with the reference but not with the ABruijn sequence (*PBcR errors* are defined analogously). Table 1 illustrates that both ABruijn and PBcR generate accurate assemblies with 25 ABruijn errors and 54 PBcR errors.

**Table 1.** Summary of putative errors for ABruijn (three polishing step) and PBcR as compared to the *E. coli K12* reference genome. All insertion and deletion errors for ABruijn and PBcR have length 1 bp with the exception of a single PBcR deletion error that has length 8 bp. The last five rows summarize putative errors for ABruijn for downsampled ECOLI dataset with reduced average coverage varying from 50X to 30X (single polishing step).

|  | Substitutions | Insertions | Deletions |
| --- | --- | --- | --- |
| *E. coli K12* | 3 | 113 | 957 |
| PBcR | 2 | 6 | 46 |
| iterative ABruijn | 0 | 7 | 18 |
| 50X | 0 | 17 | 31 |
| 45X | 0 | 26 | 47 |
| 40X | 0 | 47 | 73 |
| 35X | 0 | 86 | 130 |
| 30X | 0 | 178 | 211 |

In fact, ABruijn improves on PBcR after a single polishing step (leaving 35 errors represented by 8 insertions and 27 deletions) and further reduces the number of errors to 25 by re-aligning the reads and re-applying the polishing procedure for 3 iterations. These iterations result in the extremely low error rate of 0.0005%, which is rarely achieved even in high coverage sequencing projects with short reads. SI9: “ORF-based error-correction” describes how to further reduce the error rates by ≈ 20%.

We further estimated the accuracy of the ABruijn assembler in projects with lower coverage by downsampling the reads from ECOLI to reduce the average coverage to 50X, 45X, 40X, 35X, and 30X. For each value of coverage, we made five independent replicas and averaged the number of errors over them. Table 1 illustrates that ABruijn mantains excellent accuracy (similar to typical short read assembly projects) even in relatively low coverage projects. The lion’s share of the ABruijn errors occur in the local low-coverage regions, e.g., the accuracy in regions with coverage between 5X and 10X is rather low (on average, ≈ 1 error per 150 bps), while the accuracy in regions with coverage above 40X is extremely high. ABruijn makes ≈ 1 error per 1400, 4300, 10600, 23200, 40900, and 57600 base pairs in regions with coverage 10X-15X, 15X-20X, 20X-25X, 25X-30X, 30X-35X, and 35X-40X, respectively.

We further used ABruijn to assemble the ECOLI_*nano*_ dataset (see SI7 “Additional details on assembling Oxford Nanopore reads”). Both assembler described in Loman et al., [36] and ABruijn assembled the ECOLI_*nano*_ dataset into a single circular contig structurally concordant with the *E. coli K12* genome with error rates 1.5% and 1.1%, respectively (2475 substitutions, 9238 insertions, and 40399 deletions for ABruijn). Thus, in contrast to Pacific Biosciences technology, Oxford Nanopore technology currently has to be complemented by hybrid co-assembly with short reads to generate finished genomes [1, 30, 2,49].

While further reduction in the error rate can be achieved by machine-level processing of the signal resulting from DNA translocation [37], it is still two orders of magnitude higher that the error rate 0.008% for the down-sampled ECOLI dataset with similar 30X coverage (Table 1) and below the acceptable standards for finished genomes. Since Oxford Nanopore technology is rapidly progressing, we decided not to optimize it further using signal processing of row translocation signals.

## 5 Discussion

Since the number of bacterial genomes that are currently being sequenced exceeds the number of all other genome sequencing efforts by an order of magnitude, accurate sequencing of bacterial genomes is an important goal. Since short-read technologies typically fail to generate long contiguous assemblies (even in the case of bacterial genomes), long reads are often necessary to span repeats and to generate an accurate genome reconstruction.

Since traditional assemblers were not designed for working with error-prone reads, the common view is that OLC is the only approach capable of assembling inaccurate reads and that these reads must be error-corrected before performing the assembly [7]. We have demonstrated that both these assumptions are incorrect and that the A-Bruijn approach can be used for assembling genomes from error-prone SMRT reads. While the running time of OLC assemblers is dominated by the overlap detection step, the running time of the ABruijn assembler is dominated by the polishing step, with the assembly step itself being extremely fast (see SI10: “Running time of ABruijn”). Since this error correction step is easy to parallelize, ABruijn has the potential to become a very fast, scalable, and accurate SMRT assembler.

We have demonstrated that the ABruijn assembler works well for both Pacific Biosciences and Oxford Nanopore reads. We further introduced a new error correction approach that differs from the previously proposed approaches and generates extremely accurate genome sequences.

## 6 Acknowledgments

We are grateful to Mark Chaisson for his help with the PBcR and Quiver analysis as well as to Bahar Behsaz, Anton Korobeinikov, Mihai Pop, and Glenn Tesler for their many useful comments.

## 7. Supporting Information

### SI1: Additional details on the ABruijn algorithm

Below we provide additional details on the ABruijn algorithm.

*Bubbles in A-Bruijn graphs*. Figure S1 provides an example of a bubble in an ABruijn graph.

**Fig. S1.**
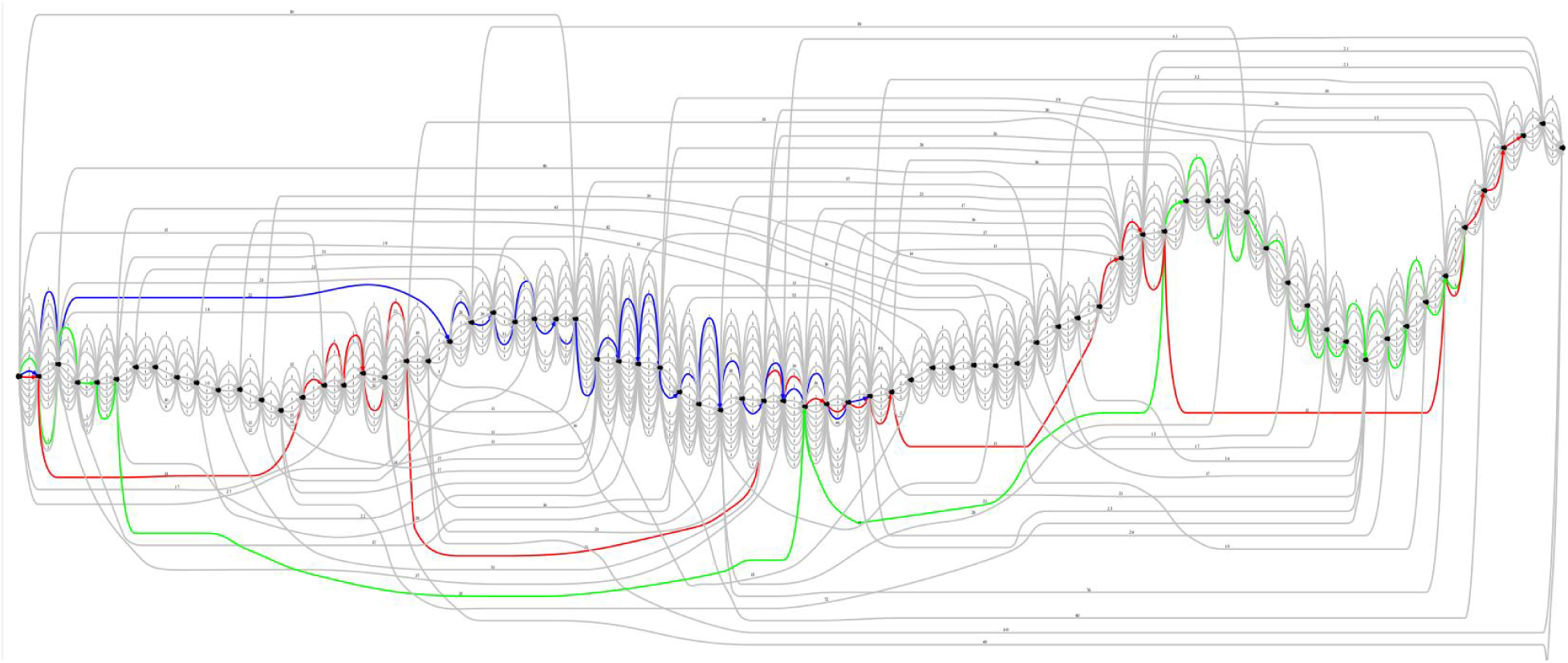
A small subgraph of the A-Bruijn graph constructed from 76 (15,8)-mers appearing in segments of 55 reads covering a short 100-nucleotide region (starting at position 2,100,000 in *E. coli* genome). Three out of 55 read-paths are highlighted in blue, red, and green.

*Fast heuristic for finding a common jump-subpath with maximum span*. We define *Predecessors*_*jump*_(*v*) as the set of all *jump*-predecessors of a vertex *v* in paths *P*_1_ and *P*_2_. A vertex *w* in *Predecessors*_*jump*_(*v*) is called *dominant* if it is not a *jump*-predecessor of any other vertex in *Predecessors*_*jump*_(*v*). If paths *P*_1_ and *P*_2_ traverse *Predecessors*_*jump*_(*v*) in the same order, then there is only one dominant vertex in *Predecessors*_*jump*_(*v*), denoted as *w*, and *span*_*jump*_(*v*) = {*span*_*jump*_(*w*) + *d*_*P*_1__ (*w*, *v*)}. To speed-up the dynamic programming algorithm based on the recurrence in the main text, ABruijn checks only the dominant vertices in *Predecessors*_*jump*_(*v*).

*Detecting chimeric reads*. BLASR alignments of reads against a genome reveal *alignable* and *unalignable* regions of each read. A read has a *low-quality end* if either its long suffix or its long prefix does not align to the genome. A read is called *chimeric* if it is formed by a concatenation of distant regions of the genome or has a long unalignable prefix or suffix (of length at least *jump*+*maxOverhang*). BLASR alignments revealed 1,983 chimeric reads in the ECOLI dataset (19% of all reads) and 19 chimeric reads in the ECOLI_*nano*_ dataset (0.3% of all reads).

The traditional way to identify a chimeric read in the de Bruijn graph framework (when the reference genome is not known) is to detect a *chimeric junction* in this read, i.e., a junction that improperly connects two parts of the genome. The existing assembly algorithms often classify a a position in the read as a chimeric junction if it is not covered by (or poorly covered by) alignments of this read with other reads. However, while this approach works for accurate reads, it needs to be modified for inaccurate reads since alignment artifacts make it difficult to identify the chimeric junctions.

Traditional de Bruijn assemblers classify a read as chimeric if one of the edges in its read-path in the assembly graph has low coverage. They further remove the chimeric reads and corresponding edges from the assembly graph (see [41] for more advanced approaches to the detection of chimeric reads). To generalize this approach to A-Bruijn graphs we need to re-define the notion of coverage for A-Bruin graphs.

**Fig. S2.**
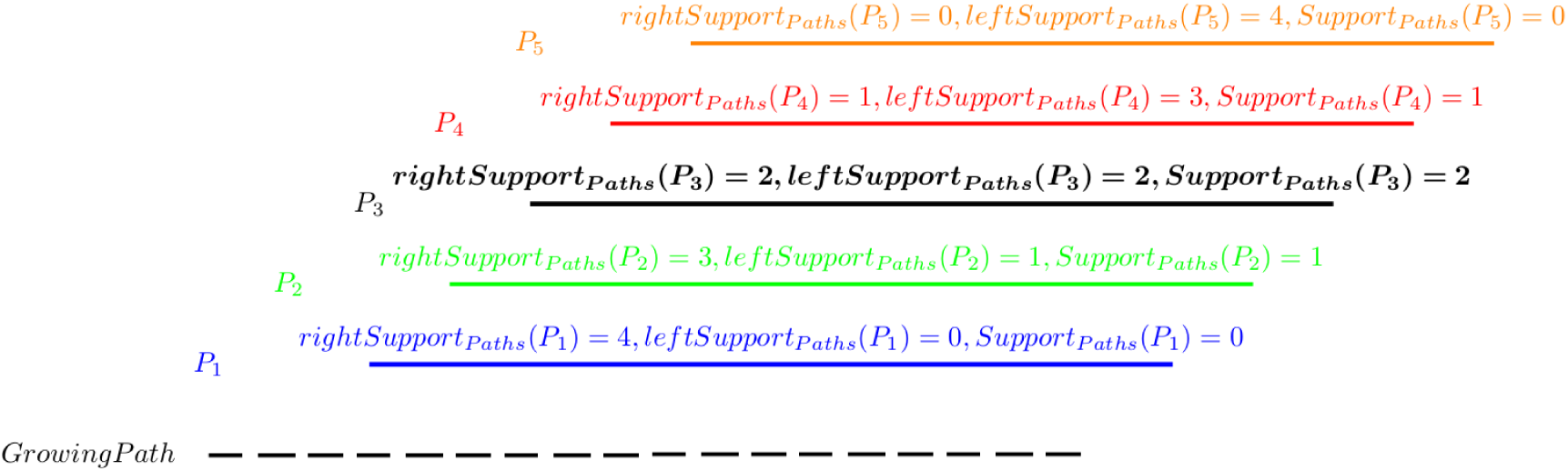
An example of *rightSupport*_*Paths*_(*P*), *leftSupport*_*Paths*_(*P*) and *Support*_*Paths*_(*P*). A *GrowingPath* and 5 paths {*P*_1_, *P*_2_, *P*_3_, *P*_4_, *P*_5_} above it (extending this path) to the right that form the set *Paths*.

An edge (*v*, *w*) in a path *P* is called *internal (strongly internal)* if the distances from *v* to the start of *P* and from *w* to the end of *P* exceed *jump* (*jump*+*maxOverhang*). Given overlapping paths *P* and *P*′, we define the *P-spread* of *P*′ as the sub-path of *P* starting and ending at the first and last vertices of *Path*_*jump*_(*P*, *P*′).

To check if a path *P* in the A-Bruijn graph is chimeric, we consider all paths *Paths* that overlap with this path and further trim non-internal edges of these paths, resulting in a set of paths that we refer to as *TrimmedPaths*. The *coverage* of an edge in path *P* is defined as the number of paths in *TrimmedPaths* whose P-spread contain this edge. A path is called *chimeric* if one of its strongly internal edges has coverage below a threshold *minCoverage* (the default value of *minCoverage* is 10% of the average coverage).

After A-Bruijn constructs the A-Bruijn graph, it runs the chimeric read detection procedure and deletes the detected chimeric reads from the A-Bruijn graph. ABruijn removes 1,931 out of 1,983 chimeric reads in the ECOLI dataset. Although 52 chimeric reads (0.5% of all reads in the ECOLI dataset) are not removed prior to constructing the genomic path in the A-Bruijn graph, no assembly errors (caused by selecting chimeric reads for path extensions) are triggered bacause the notion of a most-consistent path allows ABruijn to avoid using the chimeric reads as candidates for path extensions.

While the notion of a most-consistent path allows ABruijn to avoid selecting chimeric paths for path extensions, there is a small (0.5%) chance that it selects a chimeric read at the very first step. To make sure that ABruijn does not start from a chimeric read, we apply an additional more stringent check for chimerism to the initially selected read (i.e., increasing the value of *minCoverage* from 10% to 20% of the average coverage).

*Most-consistent paths*. Given overlapping paths *P*_1_ and *P*_2_, we say that *P*_1_ is *right-supported* by *P*_2_ if the *P*_1_-distance from the last vertex in *Path*_*jump*_(*P*_1_, *P*_2_) to the end of *P*_*1*_ is smaller than the *P*_2_-distance from the last vertex in *Path*_*jump*_(*P*_1_, *P*_2_) to the end of *P*_2_. Similarly, *P*_1_ is *left-supported* by *P*_2_ if the *P*_1_-distance from the start of *P*_1_ to the first vertex in *Path*_*jump*_(*P*_1_, *P*_2_) is smaller than the *P*_2_-distance from the start of *P*_2_ to the first vertex in *Path*_*jump*_(*P*_1_, *P*_2_). Depending on the direction of extension, the set *OverlapPaths* in the ABruijn algorithm contains all the paths in *ReadPaths* that right-support or left-support *ReadPath* (the default direction is “right”).

Given a path *P* in a set of paths *Paths*, we define *rightSupport*_*Paths*_(*P*) as the number of paths in *Paths* that right-support *P*; *leftSupport*_*Paths*_(*P*) is defined similarly. We also define *Support*_*Paths*_(*P*) as the minimum of *rightSupport*_*Paths*_(*P*) and *leftSupport*_*Paths*_(*P*).

Given a parameter *minSupport* (the default value is 2), we say that a path *P* from *Paths* is *consistent* with the set *Paths* if *Support*_*Paths*_(*P*) ≥ *minSupport*. A consistent path is *most-consistent* if it maximizes *Support*_*Paths*_(*P*).

Given a set of paths *Paths* overlapping with *ReadPath*, ABruijn selects a most-consistent path for extending *ReadPath*. It further classifies the set *Paths* as *consistent* if there exists a path *P* in this set such that nearly all other paths in *Paths* are right-supported by *P*. Note that while the simplified ABruijn pseudocode above only describes the path extension process in one direction (“right”), the ABruijn tool attempts to extend the path to the “left” if the path extension to the “right” halts.

### SI2: Choice of parameters in the ABruijn algorithm

Given parameters *k*, *t*, and *jump*, we define the following statistics (Table S1):

– *Pr*^+^(*k*, *t*, *jump*), the probability that two overlapping SMRT reads share a (*k*, *t*)-mer along a region of length *jump* in their overlap. To ensure that the notion of a common *jump*-subpath indeed detects overlapping reads, ABruijn selects parameters *k*, *t*, and *jump* in such a way that *Pr*^+^(*k*, *t*, *jump*) is large.
– *Pr*^‒^(*k*, *t*, *jump*), the probability that two regions of length *jump* from two non-overlapping SMRT reads share a (*k*, *t*)-mer. To ensure that the notion of the common *jump*-subpath does not detect non-overlapping reads, ABruijn selects parameters *k*, *t*, and *jump* in such a way that *Pr*^‒^ (*k*, *t*, *jump*) is small.

In selecting optimal parameters *k*, *t*, and *jump*, we note that the error rates in SMRT reads vary across reads and across various regions within a single read. Since the variable error rates in reads affect *Pr*^+^(*k*, *t*, *jump*) and *Pr*^‒^(*k*, *t*, *jump*), we further analyzed which parameters work the best across a range of error rates and selected the most stable ones.

ABruijn uses parameters *k* = 15, *t* = 8, and *jump* = 2000 (giving *Pr*^+^(15,8,2000) = 0.98 and *Pr*^‒^(15, 8, 2000) = 0.003) as well as *maxOverhang* = 3000 and *minOverlap* = 7000. Since most repeats in bacterial genomes have length below 7000 nt, this parameter ensures that most reads from different regions of the genome are not classified as overlapping even if they share a long repeat.

**Table S1.** The empirical estimates of *Pr*^+^(*k*, *t*, *jump*) and *Pr*^‒^ (*k*, *t*, *jump*) under different choices of parameters *k*, *t*, and *jump*. The estimates are based on statistics from 100,000 pairs of overlapping reads (to estimate *Pr*^+^ (*k*, *t*, *jump*)) and 100,000 pairs of non-overlapping reads (to estimate *Pr*^‒^ (*k*, *t*, *jump*)) from ECOLI dataset. The estimates do not significantly change for other datasets containing Pacific Biosciences reads but do change for datasets containing Oxford Nanopore reads.

| $k$ | $t$ | $jump$ | $Pr^+(k, t, jump)$ | $Pr^-(k, t, jump)$ | $k$ | $t$ | $jump$ | $Pr^+(k, t, jump)$ | $Pr^-(k, t, jump)$ |
| --- | --- | --- | --- | --- | --- | --- | --- | --- | --- |
| 15 | 7 | 2000 | 0.98 | 0.003 | 15 | 7 | 3000 | 0.99 | 0.006 |
| 15 | 8 | 2000 | 0.98 | 0.003 | 15 | 8 | 3000 | 0.99 | 0.006 |
| 15 | 9 | 2000 | 0.98 | 0.003 | 15 | 9 | 3000 | 0.99 | 0.006 |
| 15 | 10 | 2000 | 0.98 | 0.003 | 15 | 10 | 3000 | 0.99 | 0.006 |
| 17 | 6 | 2000 | 0.97 | 0.0002 | 17 | 6 | 3000 | 0.98 | 0.0005 |
| 17 | 7 | 2000 | 0.97 | 0.0002 | 17 | 7 | 3000 | 0.98 | 0.0005 |
| 17 | 8 | 2000 | 0.97 | 0.0002 | 17 | 8 | 3000 | 0.98 | 0.0005 |
| 17 | 9 | 2000 | 0.97 | 0.0002 | 17 | 9 | 3000 | 0.98 | 0.0005 |
| 19 | 5 | 2000 | 0.94 | <0.0001 | 19 | 5 | 3000 | 0.96 | 0.0001 |
| 19 | 6 | 2000 | 0.94 | <0.0001 | 19 | 6 | 3000 | 0.96 | 0.0001 |
| 19 | 7 | 2000 | 0.94 | <0.0001 | 19 | 7 | 3000 | 0.96 | 0.0001 |
| 19 | 8 | 2000 | 0.94 | <0.0001 | 19 | 8 | 3000 | 0.96 | 0.0001 |

### SI3: Additional details on analyzing necklaces

*Using the draft genome for constructing mini-alignments*. Figure S3 shows the distribution of the lengths of necklaces constructed by aligning all reads in the ECOLI dataset to the draft genome.

To evaluate how errors in the draft genome affect alignments of SMRT reads, we corrupted the reference *E. coli* genome by introducing random single-nucleotide errors at randomly chosen positions (10,000 mismatches, deletions, and insertions) and aligned all SMRT reads against the corrupted genome. A segment in the corrupted genome is called *corrupted* if it has been changed by an error and *correct* otherwise. Figure S4 shows the distribution of the local match and insertion rates (for both corrupted and correct simple 4-mers) and illustrates that 77% of all correct simple 4-mers are (0.8, 0.2)-solid. Remarkably, none of the corrupted simple 4-mers are (0.8, 0.2)-solid.

ABruijn finds all (0.8, 0.2)-solid l-mers (the default value of *l* is 10) and combines them into solid regions. It further uses the landmarks (the middle points of gold and simple 4-mers) within the solid regions as the boundaries of necklaces to ensure that single homonucleotide runs in reads do not split into two consecutive necklaces and are not adjacent to the boundaries of necklaces. This condition is important for the subsequent genome polishing step.

**Fig. S3.**
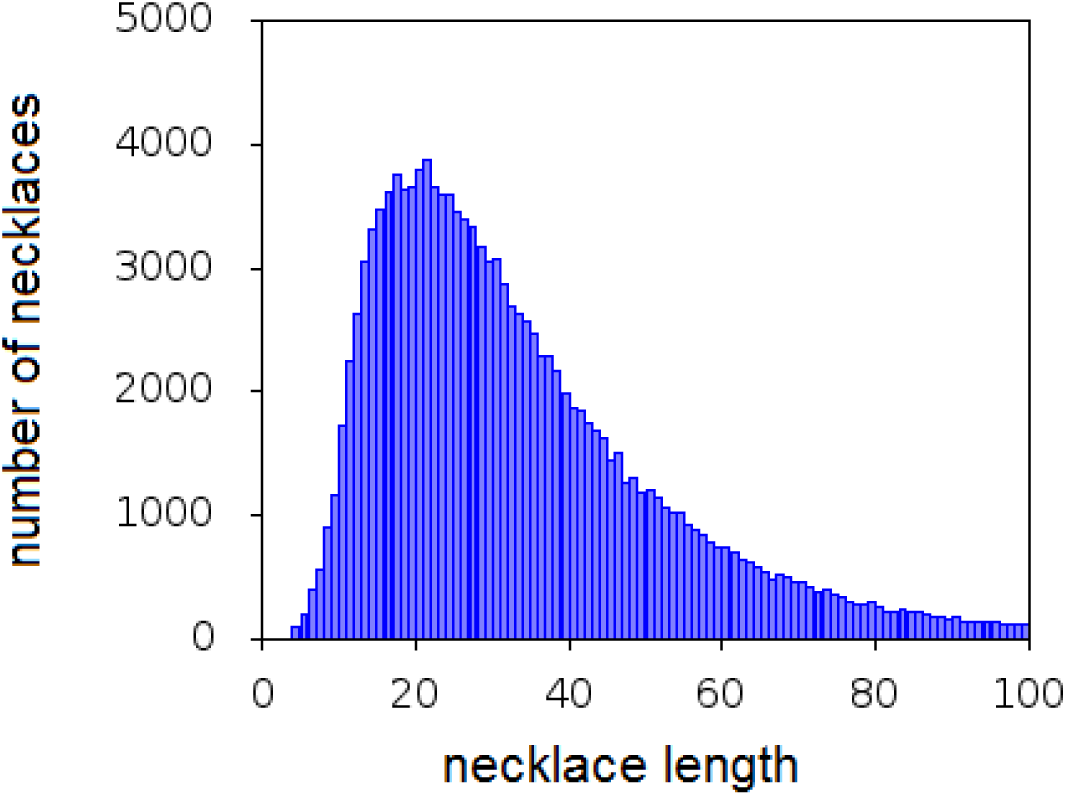
Histogram of the lengths of 135,417 necklaces formed by aligning all reads in the ECOLI dataset to the draft genome and constructing the A-Bruijn graph for that alignment. 2,914 necklaces that are longer than 100 bp are not shown.

*Selecting landmarks*. A 4-mer is called *gold* if all its nucleotides are different and *simple* if all its consecutive nucleotides are different. For example, CAGT is a gold (and simple) 4-mer, ATGA is a simple 4-mer, and GTTC is not a simple 4-mer. We further select a gold 4-mer within each solid region or, if there are no gold 4-mers, a simple 4-mer (some regions have neither gold nor simple 4-mers). We further use the middle points (i.e., a point between its 2nd and 3rd nucleotides) of selected simple 4-mers as landmarks. 135,417 out of 141,658 solid regions contain simple 4-mers resulting in 135,417 mini-alignments. ABruijn analyzes each mini-alignment and error-corrects each segment between consecutive landmarks.

*Generating necklace consensus using Partial Order Graphs*. ABruijn builds a *partial order graph (POG)* [32] for each necklace and classifies vertices in the POG as *reliable* and *unreliable*. It further constructs the path formed by reliable vertices and selects a sequence spelled by this path as the inferred consensus sequence for all reads contributing to this necklace (Figure S5). Since most necklaces are short (the average bubble length is 35 nucleotides for the ECOLI dataset), the POG construction step is fast.

We note that the Quiver algorithm [16] also uses POGs to generate consensus sequences for further polishing. However, since some details of Quiver (i.e., its scoring function) have not been published in a refereed paper yet, the differences between Quiver and ABruijn remain unclear. We thus decided to describe how ABruijn constructs and scores POGs.

Given a necklace formed by a set of segments *Segments* from reads, the pairwise alignments between the segment from the draft genome and other segments in *Segments* suffer from the fact that the draft genome is an inaccurate template for this region. While one can try to switch from this inaccurate template to another (hopefully more accurate) segment contributing to the necklace, some necklaces have no accurate segments, making such a switch problematic.

To bypass this problem, we construct a partial order graph *POG*(*Segments*), instead of relying on any single segment from one read. Each segment from *Segments* corresponds to a segment-path in the POG. The *multiplicity* of a vertex in *POG*(*Segments*) is defined as the number of segment-paths that pass through this vertex.

Given the alignment of the (unknown) reference genome to the partial order graph *POG*(*Segments*), the shared vertices between the path representing the reference genome and *POG*(*Segments*) are classified as the *reference vertices*. Intuitively, reference vertices are expected to have higher multiplicities than other vertices in *POG*(*Segments*). Indeed, our analysis of POGs constructed for the necklaces in a corrupted genome revealed that the lion’s share of reference vertices have multiplicity above |*Segments*|/2 and the lion’s share of vertices with multiplicity above |*Segments*|/2 are reference vertices.

**Fig. S4.**
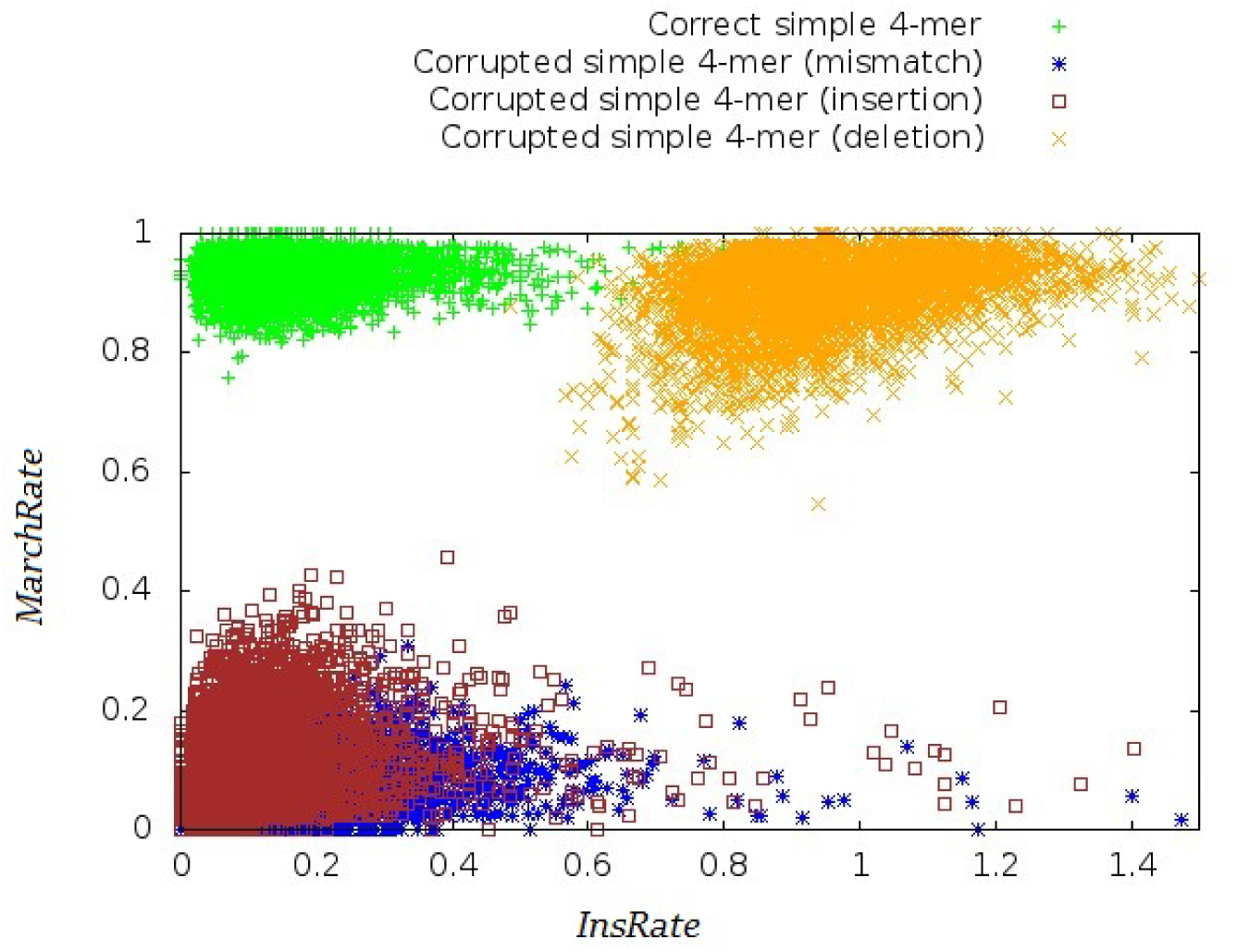
Distribution of local match and insertion rates as a 2-D plot for correct simple 4-mers (green), corrupted simple 4-mers with mismatches (blue), corrupted simple 4-mers with insertions (red) and corrupted simple 4-mers with deletions (orange).

We thus call a vertex in *POG*(*Segments*) reliable if its multiplicity exceeds |*Segments*|/2. Since the reliable vertices are totally ordered in *POG*(*Segments*), the path formed by these vertices spells out a unique sequence *consensus*(*Segments*) that we use as an initial consensus sequence of the necklace.

*Breaking long necklaces into shorter ones*. Since *POG*(*Segments*) contains pairwise alignments of all segments to *consensus*(*Segments*), we combine them into a multiple alignment and identify (0.8, 0.2)-solid *l*-mers (the default value of *l* is 10) in *consensus*(*Segments*). Using landmarks within gold and simple 4-mers within these resulting solid regions in *consensus*(*Segments*), ABruijn further decomposes *POG*(*Segments*) into shorter necklaces. This decomposition reduces the average length of necklaces from ≈35 nucleotides to ≈17 nucleotides and reduces the number of long necklaces from 2,914 to 135. Since the running time of the partial order alignment is quadratic in the total length of sequences that are being aligned, reducing the number of long bubbles results in significant reduction of the running time.

### SI4: Draft ECOLI assembly

Table S2 presents the positions of 735 reads contributing to the draft genome for the ECOLI dataset.

### SI5: Statistical analysis of errors in reads

Below we provide additional details on errors in Pacific Biosciences reads.

*Statistics of homonucleotide runs*. Table S3 presents the statistics for all read segments covering the homonucleotide runs AAAAAA and AAAAAAA. Interestingly, when we apply the statistical parameters derived from the older P5-C3 protocol to our P6-C4 ECOLI dataset, the number of ABruijn errors remains small (62 errors after a single iteration of error correction) illustrating that our probabilistic framework is not subject to over-training.

**Fig. S5.**
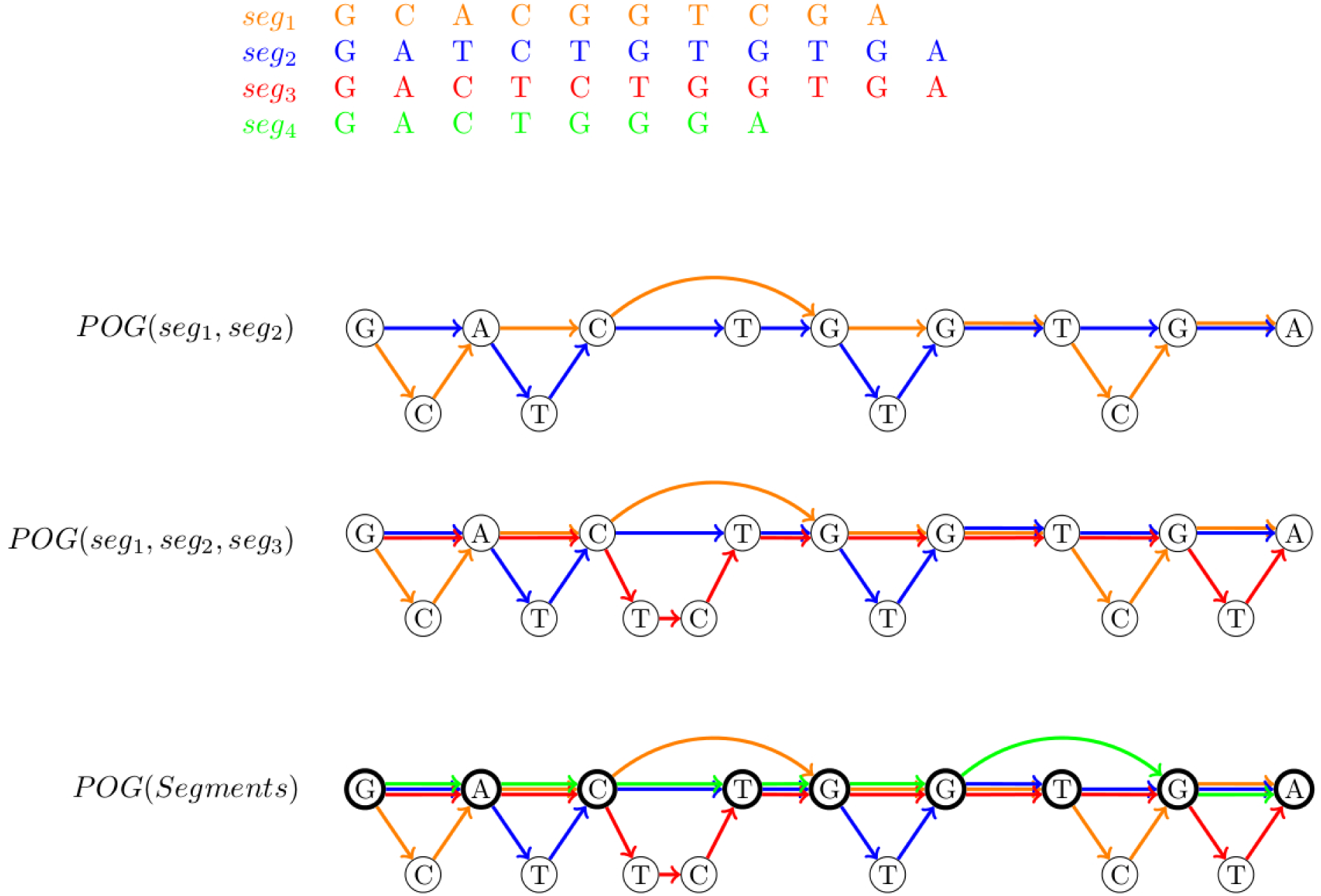
Constructing the partial order graph for *Segments* = {*seg*_1_, *seg*_2_, *seg*_3_, *seg*_4_}. Reliable vertices in *POG*(*Segments*) (shown in bold) reveal the inferred consensus sequence GACTGGTGA.

*Statistical parameters of P6-C4, P5-C3 and P4-C2 datasets*. Table S4 presents the statistical parameters of the P6-C4, P5-C3 and P4-C2 datasets.

### SI6: Differences between the genome that gave rise to the ECOLI dataset and the reference *E. coli K12* genome

Both PBcR [7, 16] and ABruijn assemblers suggest that the ECOLI dataset was derived from a strain that differs from the reference *E. coli K12* genome by a 1798 bp inversion, two insertions (776 and 180 bp), and one deletion (112 bp). To avoid confusion, we removed these regions before benchmarking ABruijn and PBcR [7, 16]. We have also clipped the PBcR assembly by ≈35 kb from the end due to a large duplication that represents a PBcR artifact when reporting the assembly of a circular genome.

PBcR assembly results in a 34,600 bp duplication at the both ends of the resulting contig which presumably represents a circularization artifact. The prefix (resp., suffix) has 54 (resp., 30) differences in this duplicated region as compared to the *E. coli K12* reference genome assembled using short Illumina reads, 5 of these differences are shared between prefix and suffix. Also the prefix (but not the suffix) contains the 1798 bp long inversion from the reference genome. Thus only counting the prefix would lead to a count of 2925 differences whereas only counting the suffix would lead to a count of 1103 differences from the reference. We include the prefix and exclude the suffix in the PBcR assembly for comparison purposes because the reads are more consistent with the presence of the 1798 bp long inversion. If we ignore the large indels and the inversion in the *E. coli K12* reference genome, then there are 45 and 59 positions with differences and ABruijn and PBcR agree on 5 of them.

### SI7: Additional details on assembling Oxford Nanopore reads

Below we provide additional details on assembling Oxford Nanopore reads.

*Modifying ABruijn to ssemble Oxford Nanopore reads*. ABruijn also assembled *ECOLI*_*nano*_ into a single circular contig structurally concordant with the *E. coli K12* genome) using the following parameters: *k* = 15, *t* = 4, and *jump* = 2500 (giving *Pr*^+^(15,4, 2500) = 0.97 and *Pr*^‒^(15,4, 2500) = 0.003). The parameters *maxOverhang* and *minOverlap* for assembling ECOLI**nano** are the same as the default parameters for ECOLI. The A-Bruijn graph was constructed from 5,997 reads, all longer than 9 kb.

To account for the specifics of Oxford Nanopore reads, we made the following changes in ABruijn as compared to assembling Pacific Biosciences reads.

– Since the ECOLI_*nano*_ dataset contains only a tiny fraction of chimeric reads, we did not remove the chimeric reads at the pre-processing stage. Instead, we relied on the notion of consistent paths to deal with chimeric reads. To avoid an accidental selection of a chimeric read at the very first step of ABruijn, we select an initial read-path that is both left-supported and right-supported by at least 3 other read-paths.
– If the set of overlapping paths with respect to a path *P* is empty (a common situation for low coverage datasets), ABruijn selects a path *P*′ that shares a common jump-subpath of maximum span with *P* and ensure that the right and left overhangs of *P* and *P*′ do not exceed *maxOverhang*.

*Constructing and error-correcting necklaces*. While the algorithm for error-correcting necklaces described in the main text is adequate for Pacific Biosciences reads, additional steps must be taken to error-correct less accurate Oxford Nanopore reads. We thus modified our polishing algorithm to account for the conservation of *k*-mers rather than the conservation of individual nucleotides (the likelihood approach described in the main text). We use *k*=5 since base-calling for Oxford Nanopore reads is based on 5-mers [37].

Given a *k*-mer in the current assembly (along with aligned reads that span this *k*-mer), we define its *support* as the number of reads for which this *k*-mer is completely conserved over the span of the alignment (i.e., the alignment contains no insertions, deletions, or substitutions over the span of the *k*-mer). We calculate the support of each *k*-mer in the current assembly and classify a *k*-mer as *weakly supported* if its support is below the *lower support threshold* and *strongly supported* if its support is above the *upper support threshold*. We observed that the lion’s share of errors are located within or nearby weakly supported *k*-mers.

For each weakly supported *k*-mer, we find the closest strongly supported *k*-mers to its left and and to its right and consider a *weak necklace* defined by reads spanning these two strongly supported *k*-mers. A mini-alignment consists of a segment of the current assembly (containing at least one weakly supported k-mer) and all read segments that are aligned to this segment. Since weak necklaces are typically very short, they often contain at least one error-free read within the span of the necklace.

ABruijn attempts to correct potential errors within each weak neaklace by analyzing alternative candidate consensus sequences. We consider two types of candidates: (i) all sequences arising from a single mutation (insertion, deletion, or substitution) in the assembled sequence, and (ii) all reads within the span of the necklace (since weak necklaces often have error-free reads).

Given a *k*-mer in a candidate sequence for a necklace, its *unaligned support* is defined as the number of read segments of that necklace that contain this *k*-mer. Unlike the previously defined (aligned) support score, the unaligned support score does not take the alignment into account in order to mitigate the problem of poorly-aligned reads within a necklace. The *candidate support score* of a candidate sequence is defined as the average value of the unaligned support scores of its *k*-mers. The algorithms selects the candidate sequence with the highest candidate support score and iterates.

*Error-correcting homonucleotide runs*. The main challenge with error-correcting less accurate Oxford nanopore reads is the deterioration of the alignment of reads against the pre-polished genome. As a result of this deterioration, the read segments that contribute to computing the consensus of a segment are not necessarily the reads that have arisen from this segment. Thus, given a segment in the pre-polished genome, we first recruit the reads that align well to this segment and only use the well-aligned reads (rather than all reads as before) for computing the likelihood. While this procedure reduces the number of reads participating in the likelihood estimation, the accuracy of the resulting consensus improves since the recruited reads are more accurate.

For each run *LZ* … *ZR* of a nucleotide *Z* in the genome flanked by the nucleotides *L* (on the left) and *R* (on the right) distinct from *Z*, we limit analysis to all reads that are well-aligned against *LZ* … *ZR*. A read is *well-aligned* against *L* … *ZR* if the flanking *L* (*R*) nucleotide forms either a match with the read or is aligned against a nucleotide *Z* in the read. We represent the segment of the well-aligned read simply as the count of the nucleotides contained in the run and the count of all other nucleotides (see Figure S7).

**Fig. S6.**
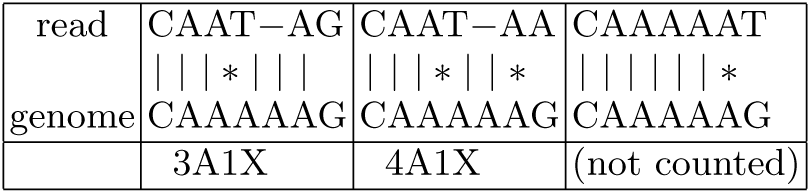
Well-aligned reads (the first two examples) and a poorly aligned read (the last example). The well-aligned reads are represented as 3A1X and 4A1X in the likelihood estimate (X stands for an arbitrary nucleotide.

After generating all well-aligned read-segments for all homonucleotide runs, we compute the conditional probability that a homonucleotide run of a given length results in a read-segment of a given type (e.g., *Pr*(read-segment=4A1X | genome-segment=5A)=0.0585) as well as the probability of each homonucleotide run (e.g., *Pr*(genome-segment=5A)=0.0075) (see Table S5). We further use the resulting probabilities to compute the likelihood function for all well-aligned reads in each necklace (frequencies below a threshold 0.001 are ignored). Our analysis revealed tha these probabilities hardly change when one changes the dataset of reads, coverage, or the reference genome. Given a read-segment from a well-aligned read, we define *Pr*(read-segment, run-length) = *Pr* (read-segment—run-length) · *Pr*(run-length)

Given a set of read-segments *Segments* from all well-aligned reads, we select the run-length of a homonucleotide run in a necklace as the run-length that maximizes the formula above.

For example, if *Segments*={3A, 4A, 4A1X, 4A1X, 5A}: *Pr*(*Segments*, 4*A*) = *Pr* (*Segments*|4*A*) · *Pr*(4*A*) = 0.3512 · 0.3593 · 0.035 · 0.035 · 0.0153 · 0.0204 = 4.8^‒8^

*Pr* (*Segments*, 5*A*) = *Pr* (*Segments*|5*A*) · *Pr*(5*A*) = 0.2559 · 0.3106 · 0.0585 · 0.0585 · 0.1259 · 0.0075 = 2.7^‒7^

Since *Pr*(5*A*|*Segments*) > *Pr*(4*A*|*Segments*), we select AAAAA over AAAA as the length of the homonucleotide run.

### SI8: Assembling *Xanthomonas* genomes

Since HGAP 2.0 failed to assemble the BLS256 genome, Booher et al., 2015 [10] developed a special PBS algorithm for “local *tal* gene assembly and applied Minimo assembler [57] to address this deficiency in HGAP. They further proposed a workflow that first launches PBS and uses the resulting local *tal* gene assemblies as seeds for a further HGAP assembly with custom adjustment of parameters in HGAP/Celera workflows (this workflow was used for assembling the PXO99A genome as well). While HGAP 3.0 resulted in an improved assembly of BLS256 (as compared to HGAP 2.0), Booher et al., 2015 [10] commented that the PBS algorithm is still required for assembling other *Xanthomonas* genomes. We further comment that PBS represents a customized assembler for *tal* genes that is not designed to work with other types of tandem repeats. Thus, development of a general SMRT assembly tool that accurately reconstruct arbitrary tandem repeats remains an open problem.

Since BLS256 and PXO99A have various high-multiplicity repeats, *k*-mers from these repeats have very high frequencies. Since ABruijn excludes high-frequency *k*-mers from the construction of the A-Bruijn graph (Figure 2), *k*-mers from TAL and IS repeats in *Xanthomonas* genomes become invisible in the A-Bruijn graph. This makes finding common *jump*-paths in reads containing TAL and IS repeats problematic and makes it difficult to identify overlapping reads from these regions. To address this challenge, we modified ABruijn for assembling tandem repeats with high copy numbers as follows.

The modified ABruijn assembler does not remove all *k*-mers with high frequencies exceeding *c* · *t* (the default *c*=3) from the set of solid strings in the construction of the A-Bruijn graph. Instead, it down-samples all high frequency k-mers (with frequencies exceeding *c* · *t*). This down-sampling randomly selects at most *c* · *t* occurrences of a frequent *k*-mer in reads (rather than excluding all frequent *k*-mers) when forming read-paths, thus preventing fragmentation in the resulting assembly caused by overly-aggressive removal of high-frequency *k*-mers in TAL and IS repeats. An additional challenge is the potential misalignment of SMRT reads spanning tandem TAL repeats in the A-Bruijn graph (which may erroneously align the i-th and *j*-th copies of a tandem TAL repeat for *i* ≠ *j*). To minimize the effects of such misalignments, we added an additional constraint on vertices in common *jump*-subpaths by requiring that |*d*_*P*_1__ (*v*_*i*_, *v*_*i*+1_) – *d*_*P*_2__ (*v*_*i*_, *v*_*i*+1_)| ≤ *jump*/2 for all adjacent vertices *v*_*i*_ and *v*_*i*+1_ in all common *jump*-subpaths between read-paths *P*_1_ and *P*_2_.

We launched ABruijn with the following parameters to assemble the BLS256 and PXO99A datasets: *k*=15, *t*=8, *jump*=2000, *maxOverhang*=1500 and *minOverlap*=8000. ABruijn assembled the BLS256 genome into a single circular contig structurally concordant with the BLS256 reference genome. It also assembled the PXO99A genome into a single circular contig structurally concordant with the PXO99A reference genome but, similarly to the initial assembly in Booher et al., 2015 [10], it collapsed a 212 kb tandem repeat.

### SI9: ORF-based error-correction

While the likelihood-based approaches to error-correction (described in the main text) corrects the lion’s share of errors in the draft genomes, some errors remain uncorrected, particularly with respect to the errors in estimating the lengths of homonucleotide runs. We thus complement the likelihood-based approaches with a new *ORF-based error-correction* approach that analyses Open Reading Frames (ORFs).

Note that while the average length of a protein-coding gene in most bacterial genomes exceeds 800 [11], the average ORF length in a randomly generated string of nucleotide is only 64. Thus, every error that represents an indel within a gene (a frameshift) may introduce a premature stop codon and has the potential to significantly reduce the length of the ORF corresponding to this gene.

If we are deciding between two alternative lengths of a homonucleotide run within a gene (correct and incorrect), the correct choice results in an ORF that corresponds to the gene length while the incorrect choice results in a frameshift that may introduce a premature stop codon. Such frameshift mutations usually shorten the length of the longest ORF that spans over the homonucleotide run with incorrectly defined length.

Given a position in the genome, we compute its *ORF-length* as the maximum length of all six ORFs covering this position. If the genome is assembled without errors then ORF-lengths are large for most positions that belong to genes. Since genes typically cover over 85% of bacterial genomes, most positions in the entire genome have large ORF-lengths. However, if a genome is assembled with errors, the ORF-lengths for positions with indels are typically smaller than the ORF-length of this position in the error-free genome (see Figure S7).

Since in some cases, the likelihood values for alternative choices for the length of a homonucleotide run are nearly the same, we develop an additional decision rule that analyzes the ORF-lengths between two alternatives and gives preference to the choice that results in a significantly longer ORF-length.

Given two candidate lengths of a homonucleotide run with a small difference in their homonucleotide likelihood score (smaller than a threshold *Δ*), we compute the difference between their ORF-lengths and select the candidate with larger ORF-length if the difference between ORF-lengths exceeds a threshold (the default value is 128 bp). If the difference between the ORF-lengths is smaller than the threshold, we retain the length of the run that maximizes the homonucleotide likelihood score described in the main text.

### SI10: Running time of ABruijn

The construction of the A-Bruijn graph (using *k*-mer counting program DSK [50] and naive *k*-mer indexing) and finding a genomic path in this graph together take under 30 min using modern 8-core desktop computer with 32 Gb memory (for all genomes we analyzed). We estimate that using a fast rather than naive *k*-mer indexing algorithm would significantly reduce the time for constructing the draft genome.

Since the ABruijn assembly step is very fast, its running time is dominated by the polishing step that is currently implemented in Python; it takes about 6 hours on the ECOLI dataset (similar to the time taken by Quiver error correction step in PBcR). We estimate that the running time of the polishing step wil be reduced by an order of magnitude after we complete the transition from Python to C++.

ABruijn assembler is freely available from https://sites.google.com/site/abruijngraph

**Table S2.**
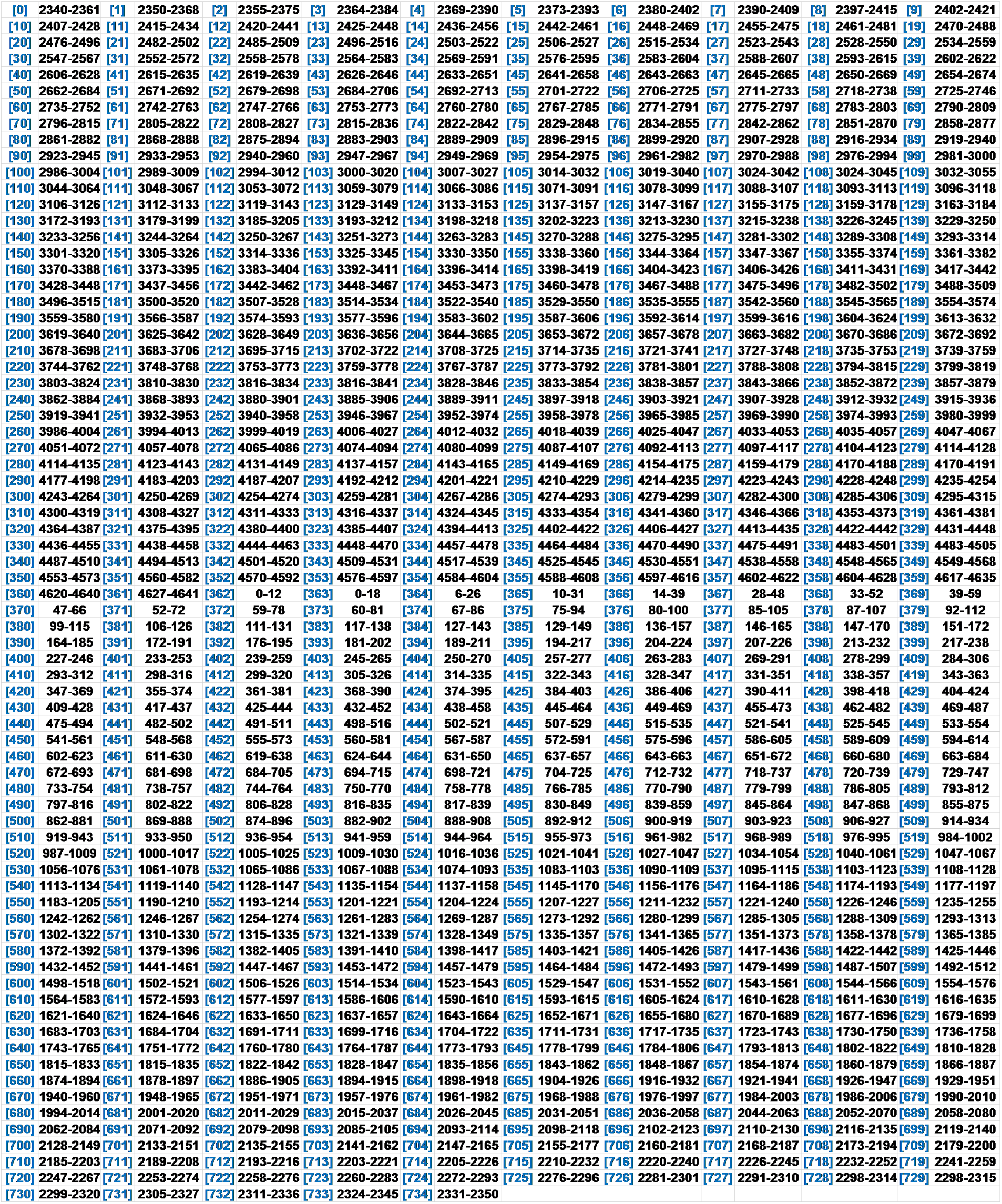
Summary of the positions of 735 reads contributing to the draft genome for the ECOLI dataset. The order of reads (from 0 to 734) corresponds to the order that ABruijn used to execute the path extension paradigm. Each read is represented by the starting and ending position of its alignment to the reference genome (rounded to the nearest 1000). For example, read 0 is aligned to positions between 2,340 kb and 2,361 kb in the reference genome. Note that the length of a read may be longer than the span of the alignment since many reads have low-quality ends that do not align to the genome.

**Table S3.** The counts and frequencies of segments from reads spanning 6-nucleotide runs AAAAAA (left) and 7-nucleotide runs AAAAAAA (right) in *E. coli* genome. Only the combinations with frequencies exceeding 0.001 are shown.

| AAAAAA | Count | Frequency | AAAAAAA | Count | Frequency |
| --- | --- | --- | --- | --- | --- |
| 6A | 42522 | 0.473 | 6A | 3455 | 0.154 |
| 5A | 13970 | 0.155 | 5A | 1104 | 0.049 |
| 7A | 8997 | 0.100 | 7A | 9360 | 0.418 |
| 4A | 3538 | 0.039 | 8A | 2339 | 0.104 |
| 6A1T | 2139 | 0.024 | 7A1T | 569 | 0.025 |
| 6A1G | 2075 | 0.023 | 7A1G | 545 | 0.024 |
| 5A1C | 1824 | 0.020 | 9A | 510 | 0.023 |
| 6A1C | 1785 | 0.020 | 6A1C | 486 | 0.022 |
| 8A | 1512 | 0.017 | 7A1C | 401 | 0.018 |
| 5A1T | 1265 | 0.014 | 6A1T | 372 | 0.017 |
| 5A1G | 1169 | 0.013 | 6A1G | 302 | 0.013 |
| 3A | 845 | 0.009 | 4A | 300 | 0.013 |
| 4A1C | 498 | 0.006 | 5A1C | 150 | 0.007 |
| 7A1G | 426 | 0.005 | 8A1G | 133 | 0.006 |
| 7A1T | 407 | 0.005 | 10A | 114 | 0.005 |
| 7A1C | 316 | 0.004 | 8A1T | 111 | 0.005 |
| 6A1T1G | 261 | 0.003 | 8A1C | 100 | 0.004 |
| 9A | 259 | 0.003 | 5A1T | 91 | 0.004 |
| 4A1T | 238 | 0.003 | 5A1G | 76 | 0.003 |
| 4A1G | 214 | 0.002 | 3A | 74 | 0.003 |
| 6A1C1G | 210 | 0.002 | 7A2G | 68 | 0.003 |
| 6A2C | 197 | 0.002 | 7A1T1G | 65 | 0.003 |
| 6A2G | 196 | 0.002 | 7A1C1G | 65 | 0.003 |
| 6A2T | 183 | 0.002 | 6A2C | 48 | 0.002 |
| 6A1C1T | 175 | 0.002 | 7A1C1T | 46 | 0.002 |
| 2A | 172 | 0.002 | 7A2T | 45 | 0.002 |
| 5A2C | 151 | 0.002 | 6A1C1G | 40 | 0.002 |
| 3A1C | 131 | 0.001 | 6A1C1T | 39 | 0.002 |
| 5A1T1G | 114 | 0.001 | 7A2C | 38 | 0.002 |
| 5A1C1T | 113 | 0.001 | 6A1T1G | 36 | 0.002 |
| 4A2C | 110 | 0.001 | 4A1C | 35 | 0.002 |
| 5A1C1G | 107 | 0.001 | 5A2C | 32 | 0.001 |
| 5A2G | 101 | 0.001 | 9A1G | 32 | 0.001 |
| 1A | 94 | 0.001 | 9A1T | 28 | 0.001 |
|  |  |  | 6A2G | 27 | 0.001 |
|  |  |  | 2A | 25 | 0.001 |
|  |  |  | 5A1C1T | 24 | 0.001 |
|  |  |  | 8A1T1G | 23 | 0.001 |

**Table S4.** Comparison of statistical parameters of the P6-C4 protocol with the statistical parameters of the older P5-C3 and P4-C2 protocol (derived from the P5-C3 and P4-C2 SMRT datasets in [27]).

|  | P6-C4 | P5-C3 | P4-C2 |
| --- | --- | --- | --- |
| *Match(A)* | 0.958 | 0.962 | 0.953 |
| *Match(C)* | 0.944 | 0.935 | 0.946 |
| *Match(G)* | 0.950 | 0.946 | 0.920 |
| *Match(T)* | 0.956 | 0.958 | 0.936 |
| *Sub(A → C)* | 0.005 | 0.004 | 0.006 |
| *Sub(A → G)* | 0.002 | 0.002 | 0.002 |
| *Sub(A → T)* | 0.002 | 0.002 | 0.003 |
| *Sub(C → A)* | 0.008 | 0.012 | 0.010 |
| *Sub(C → G)* | 0.004 | 0.006 | 0.002 |
| *Sub(C → T)* | 0.004 | 0.004 | 0.003 |
| *Sub(G → A)* | 0.004 | 0.004 | 0.006 |
| *Sub(G → C)* | 0.003 | 0.003 | 0.006 |
| *Sub(G → T)* | 0.004 | 0.004 | 0.006 |
| *Sub(T → A)* | 0.004 | 0.004 | 0.006 |
| *Sub(T → C)* | 0.003 | 0.002 | 0.007 |
| *Sub(T → G)* | 0.004 | 0.003 | 0.004 |
| *Del(A)* | 0.032 | 0.030 | 0.036 |
| *Del(C)* | 0.041 | 0.043 | 0.038 |
| *Del(G)* | 0.039 | 0.043 | 0.062 |
| *Del(T)* | 0.033 | 0.033 | 0.047 |
| *Ins(A)* | 0.027 | 0.028 | 0.024 |
| *Ins(C)* | 0.019 | 0.016 | 0.031 |
| *Ins(G)* | 0.022 | 0.024 | 0.016 |
| *Ins(T)* | 0.021 | 0.020 | 0.016 |
| *NoIns* | 0.912 | 0.912 | 0.913 |

**Table S5.**
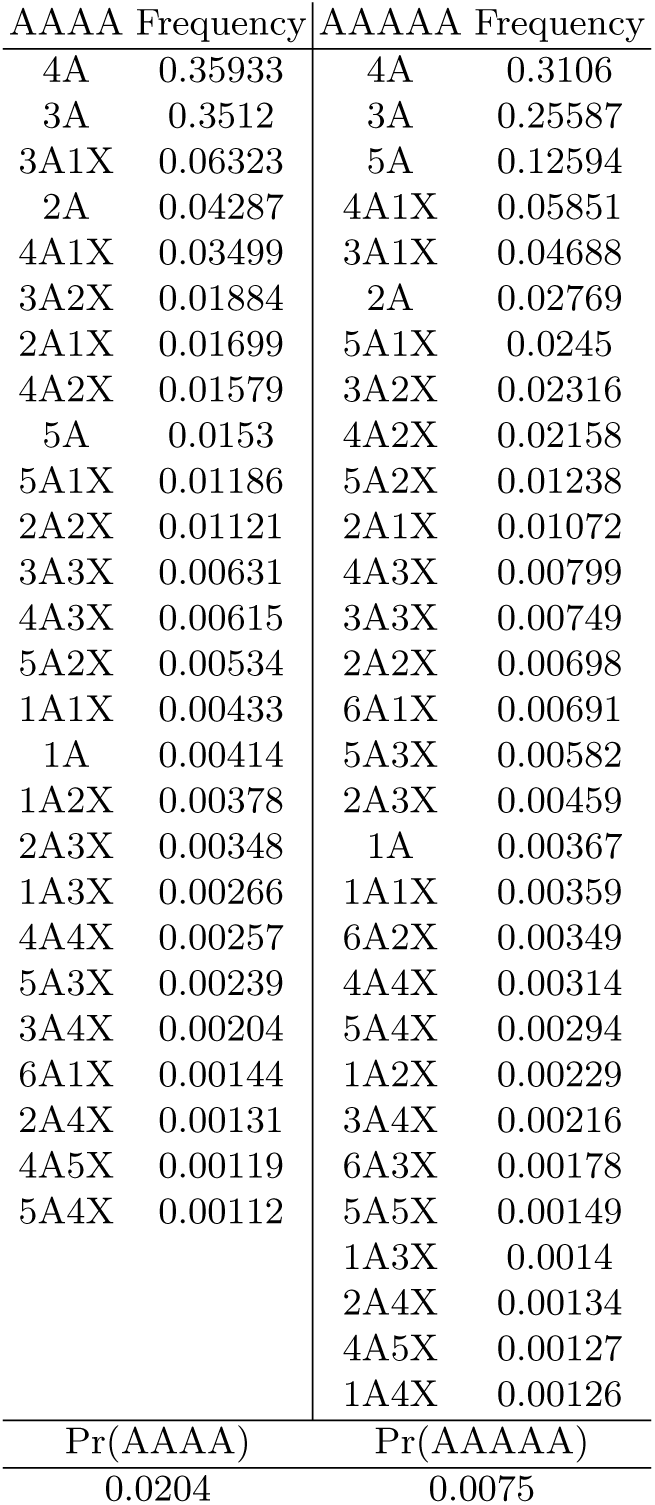
The counts and frequencies of segments from Oxford Nanopore reads spanning 4-nucleotide runs AAAA (left) and 5-nucleotide runs AAAAA (right) in *E. coli* genome. Only the combinations with frequencies exceeding 0.001 are shown. X stands for an arbitrary nucleotide.

**Fig. S7.**
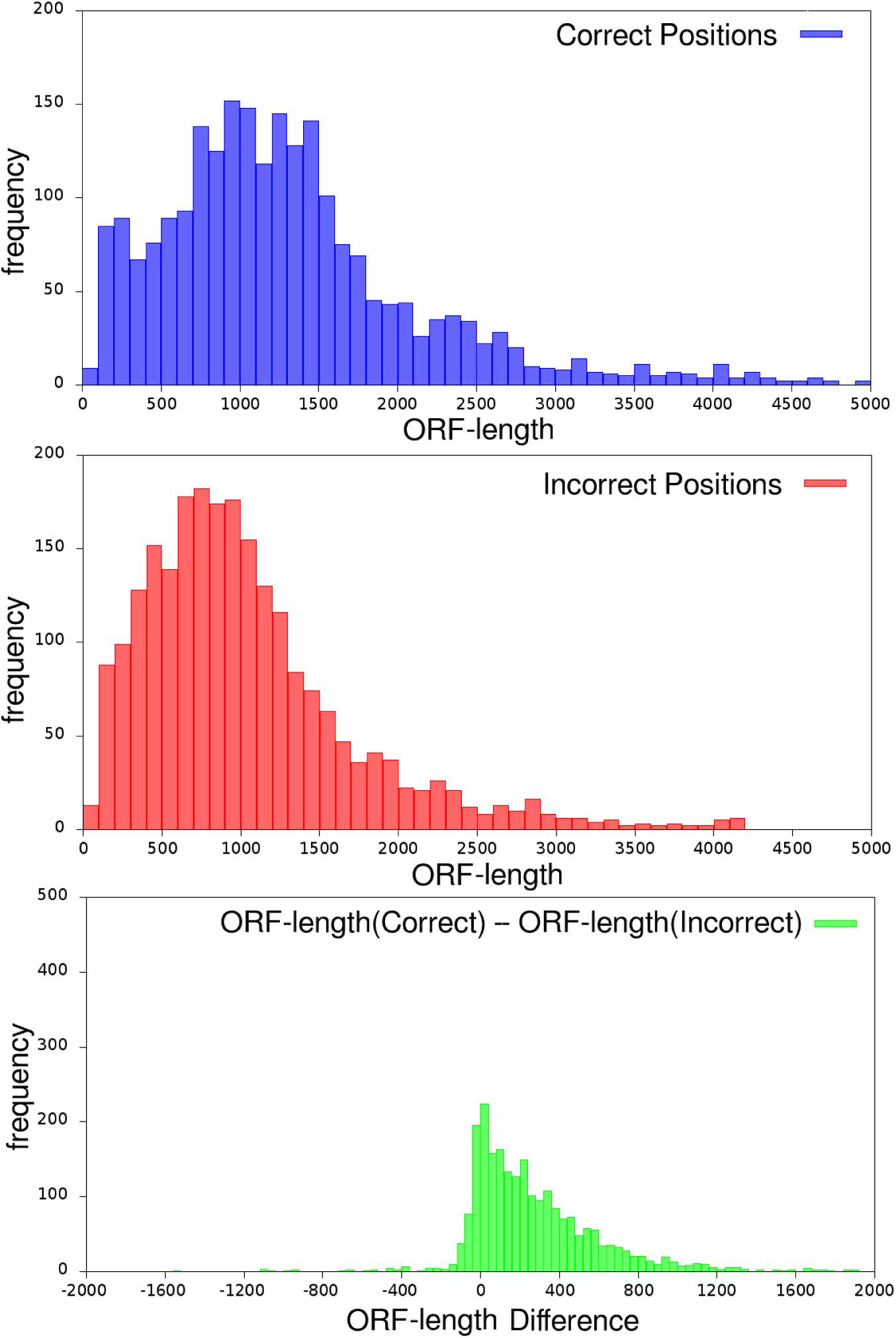
Distribution of ORF-lengths for correct positions in the error-free *E. coli* genome (top) and incorrect positions in the error-prone *E. coli* genome (middle), and the difference between the ORF-lengths of corresponding correct and incorrect positions (bottom). The error-prone *E. coli* genome was generated by deleting or inserting a single (randomly chosen) nucleotide with probability 0.0005 at each position. The vast majority of indels in the error-prone genome result in a significant reduction of ORF lengths. On average, there is a 276 nucleotide reduction in the ORF-length for the error-prone genome.

